# A roadmap to mammalian oral microbiome evolution with dental calculus

**DOI:** 10.1101/596791

**Authors:** Jaelle C. Brealey, Henrique G. Leitão, Tom van der Valk, Wenbo Xu, Katia Bougiouri, Love Dalén, Katerina Guschanski

## Abstract

Animals and their associated microbiomes share a long evolutionary history, influenced by a complex interplay between extrinsic environmental and intrinsic host factors. However, we know little about microbiome responses to long-lasting environmental and host-centred processes, which require studying microbiome changes through time. Here, we apply a temporal metagenomics approach to dental calculus, the calcified oral microbial biofilm. We establish dental calculus as a valuable tool for the study of host microbiome evolution by characterising the taxonomic and functional composition of the oral microbiome in a variety of wild mammals. We detect oral pathogens in individuals with evidence of oral disease, assemble near-complete bacterial genomes from historical specimens, characterise antibiotic resistance genes even before the advent of industrial antibiotic production, reconstruct components of the host diet and recover host genetic profiles. Our work demonstrates how dental calculus can be used in the future to study the evolution of oral microbiomes and pathogens, and the impact of anthropogenic changes on wildlife and the environment.

## Main

Hosts and their associated microbiomes share a long evolutionary history, influenced by a complex interplay between host genetics and the environment^1–4^. On short temporal scales, experimental work in model organisms has provided important insights into a multitude of factors shaping the host-associated microbial community, including the effect of diet, reproductive status and rapid environmental change^5–7^. However, there is limited understanding of how the host-associated microbiome is influenced by population-level extrinsic and intrinsic processes that span multiple host generations, such as long-term environmental and host demographic changes. Similar to fossilised plant and animal remains, microbial fossils can offer a unique opportunity to study microbiome evolution and quantify how complex microbial communities and their individual members change through time. Unfortunately, host-associated microbiomes rarely persist after the host’s death. An accessible and well-preserved exception is dental calculus, the calcified form of the microbial biofilm that forms on mammalian teeth^8^. Besides representing the oral microbial community, dental calculus also captures invading pathogens, host cells and dietary biomolecules, thus providing the opportunity for integrative investigations of microbial, dietary and host genetic factors from the same source material through sequencing of preserved DNA^9–12^. Dental calculus undergoes periodic mineralisation during an individual’s life, which reduces external contamination and facilitates DNA preservation through time^8^.

To date, dental calculus has been studied in humans, where DNA sequencing has revealed shifts in oral microbiome composition associated with cultural transitions and allowed tracking of host-pathogen co-evolution through time^9–11^. Many other mammals produce dental calculus, offering the opportunity to study the link between evolutionary, ecological and demographic factors, and microbiome evolution. Yet, this rich source of information remains virtually unexplored. Dental calculus is readily available from museum-preserved and archaeological specimens. Natural history collections have been extensively used to study the effects of human-driven changes in animal populations over the last few hundred years, such as population declines, range shifts and introduction of pollutants^13–16^. It may also be possible to infer the effects of these processes on host-associated microbiomes and the potential of microbiomes to confer adaptations to changing environments^17^. To establish dental calculus as a standard research tool for the study of host-associated microbiome evolution in diverse mammalian species, we used DNA sequencing to characterise the historical dental calculus microbiome of three evolutionarily distant mammalian species with distinct ecology, diet and physiology: European reindeer (*Rangifer tarandus*), Scandinavian brown bear (*Ursus arctos*), and eastern gorilla (*Gorilla beringei*). Reindeer are group-living ruminant herbivores with a multigastric digestive system and specialised hypsodont molars adapted to an abrasive, fibrous diet. Brown bears are solitary omnivores with brachydont molars more adapted for a partially carnivorous diet. Gorillas are group-living folivores and specialised hindgut fermenters. The close evolutionary relationship between gorillas and humans (the major source of the microbial reference databases used for microbiome taxonomic characterisation) and the previous successful reconstruction of the chimpanzee oral microbiome from dental calculus^11^ prompted us to include gorillas to aid characterisation of previously unexplored microbiomes from the other two host species. We outline strategies to overcome the challenges of working with historical microbial DNA from non-model host species, including contamination issues and reference database biases, and demonstrate that a wealth of evolutionary, ecological and conservation-relevant information can be obtained from historical dental calculus samples of diverse host species.

## Results

### Oral microbiome signature can be successfully recovered from dental calculus of non-human mammals

DNA extraction, Illumina shotgun sequencing and metagenomics analyses were carried out on dental calculus collected from five reindeer (including forest, mountain and high arctic Svalbard ecotypes), six brown bears (from western and eastern Europe) and two eastern gorillas (one Grauer’s and one mountain gorilla) dating from 1861 to 1961 (Supplementary Table 1). Contamination of samples with modern DNA is a major problem faced by all historical genomic and metagenomic studies^18^. We used a combination of laboratory and bioinformatics procedures to quantify and reduce contamination. In the laboratory, we tested three surface decontamination treatments on the two gorilla dental calculus samples: UV exposure, a wash in EDTA-containing buffer, and no surface decontamination as a control. Following microbial taxonomic assignment with Kraken2^19^, we determined that samples decontaminated with the EDTA wash contained the highest proportion of oral taxa (Supplementary Figure 1), and they were therefore retained for downstream analyses. We employed a multi-step bioinformatics approach, which relied on the availability of negative controls and the hallmarks of ancient DNA to identify and filter out contaminant taxa. More specifically, we flagged taxa as contaminants if they were present in the negative controls from the extraction and library preparation stages^20^, had higher relative abundance in low biomass samples^21^, had predominantly long DNA fragments (Supplementary Figure 2) and lacked the typical post-mortem deamination patterns^18^ (Supplementary Figure 3).

To determine the proportion of microbial communities in our samples that could be assigned to either well-characterised oral microbiome taxa or putative contaminating sources, we used the Bayesian classification tool SourceTracker^22^. While there was substantial variation between samples, an oral microbiome signature was present in dental calculus samples from all three host species (Figure 1a). Generally, the oral microbiome signature was highest in the two gorilla individuals (Supplementary Figure 4), likely the result of improved taxonomic classification due to the close phylogenetic relationship between gorillas and the source of the oral microbiome database (humans). Human skin and laboratory reagents were the most common contamination sources, and were particularly abundant in specimens that did not undergo surface decontamination before DNA extraction (Supplementary Figure 5). One such bear sample (Ua6) contained high levels of contaminants (>70%) and no detectable oral microbiome signature, and was therefore excluded from all microbial analyses.

**Figure 1.**
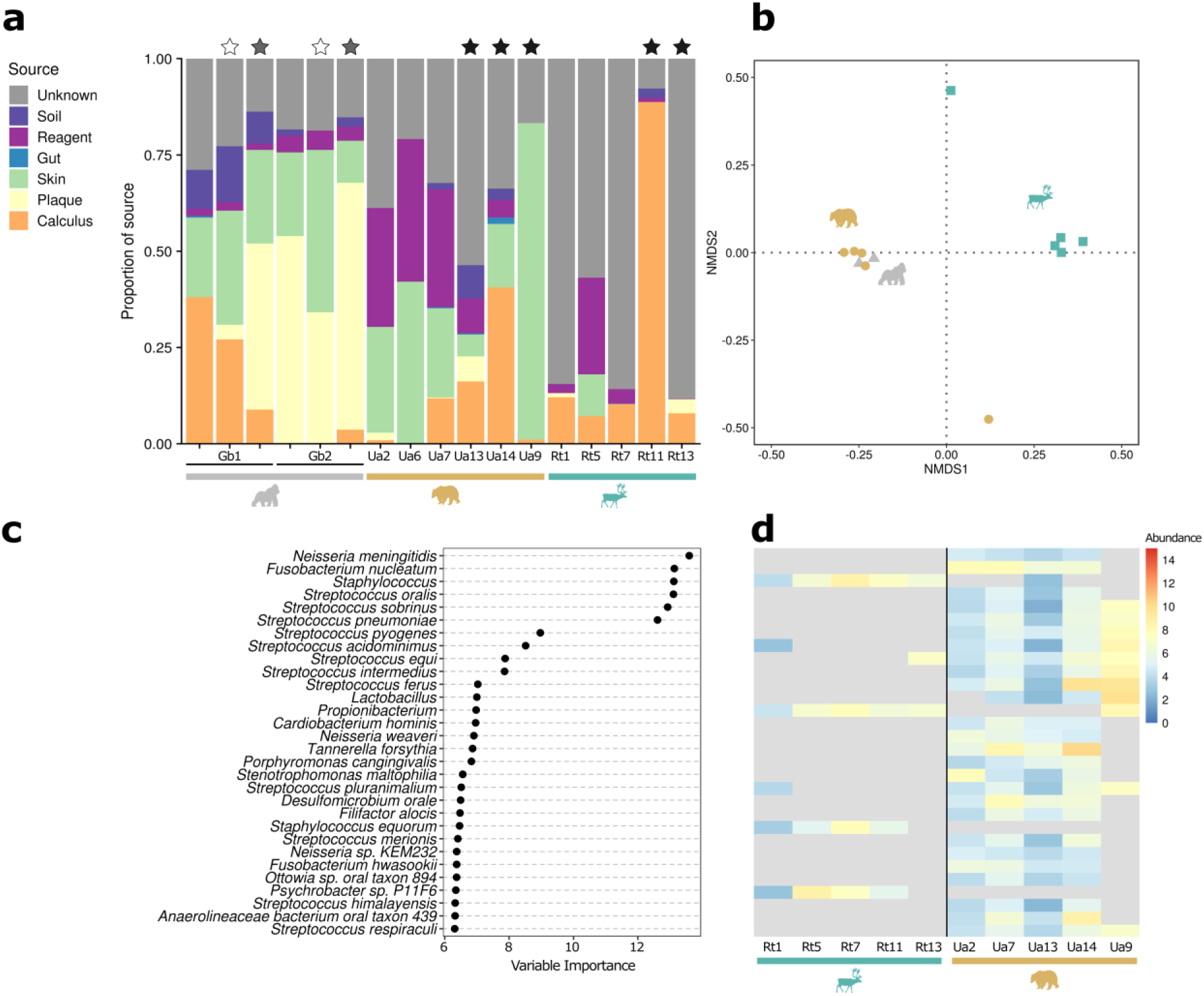
Dental calculus of non-human mammals shows an oral microbiome signature and contains host-specific taxa. **a)** Proportions of source contributions to the microbial communities (identified taxonomically at the species and genus level) contained in the dental calculus samples estimated by SourceTracker. Stars above bars indicate samples in which surface decontamination was performed before DNA extraction (white: UV only, grey: EDTA wash only, black: UV followed by EDTA wash). Note that each gorilla sample was processed in three different ways to determine the most efficient decontamination strategy (see Methods). **b)** NMDS ordination on Jaccard presence/absence distance matrix of microbial taxa per sample, coloured by host species. **c)** Random forest variable importance plot of the 30 most discriminatory taxa comparing bear and reindeer samples, based on presence/absence data after contamination filtering. **d)** CLR normalised abundance of the top 30 taxa in **(c)** in the bear and reindeer samples. Taxa that were not detected in a sample are coloured grey.

Host species had distinct oral microbiome composition (Figure 1b; for abundance-based comparison see Supplementary Figure 6 and Supplementary Table 2). Approximately 30% of the variation between samples could be explained by host species (*p*=0.006) and < 10% each by surface decontamination, sequencing depth and abundance of human reads as measure of human DNA contamination (Supplementary Table 2). Microbial taxa that showed the greatest host species specificity (i.e. were only present in a single host species) were generally associated with either the mammalian oral microbiome or other mammalian body sites (Supplementary Table 3). To independently verify the host species-specific differences between reindeer and bear (gorillas were excluded due to the low sample size of 2), we used a random forests classifier on presence/absence data. The correct host species could be assigned in 90.0% of cases, and the most important taxa for determining the host species generally included oral taxa, such as *Streptococcus* species, which were highly abundant in the bears but almost entirely absent in reindeer (Figure 1c-d). Reindeer individuals also had lower microbial diversity than bears and gorillas (Supplementary Figure 7, Shannon diversity index median ± interquartile range: 1.777 ± 0.288 (reindeer) vs. 3.794 ± 0.014 (gorilla) vs. 3.836 ± 0.602 (bear), p=0.003). This low alpha diversity of the reindeer samples may reflect poor representation of reindeer oral taxa in the reference database, rather than true reduced diversity.

### Oral pathogens in wild animals and functional repertoire of the mammalian oral microbiome

While oral diseases are often thought of as an affliction of human dietary practices^23^, oral pathologies have been observed in animals^24,25^, including wild North American black bears^26^ and captive brown bears^27^. One of the brown bear specimens in our study (Ua9) was sampled from a caries lesion (Figure 2a). The most abundant microbial taxa in Ua9 included members of the *Lactobacillus casei* group (*L. casei*, *L. paracasei*, *L. rhamnosus* and *L. zeae*) and mutans streptococci (such as the closely related *Streptococcus mutans* and *S. ratti*^28^), species that have been associated with caries lesions in humans^29,30^ (Figure 2b). Species in the *L. casei* group in particular have been associated with active caries lesions in humans^29,30^. One additional bear individual had signs of caries (Ua7), but was sampled from a healthy tooth, rather than a caries lesion, and its microbiome taxonomically appeared more similar to the other bear samples without macroscopic signs of oral disease (Figure 2b). The biogeography of the oral microbiome in humans has been found to be site-specific^31^, suggesting that even within a diseased individual, the oral microbiome could differ between healthy and carious sites. Cariogenic bacteria including mutans streptococci and *S. mitis* group species were identified in most bear samples without signs of oral disease, as were bacterial species associated with the periodontal ‘red complex’ pathogens (*Porphyromonas gingivalis*, *Treponema denticola*, and *Tannerella forsythia*)^32^. However, these samples lacked members of the *L. casei* group. Human studies suggest that dental caries is a complex disease that depends more on the environment of the oral cavity and the functions performed by the microorganisms, than on the specific taxonomic composition^33^. The same potentially pathogenic taxa were also found in gorilla and some reindeer samples, suggesting that they represent members of the healthy mammalian oral cavity.

**Figure 2.**
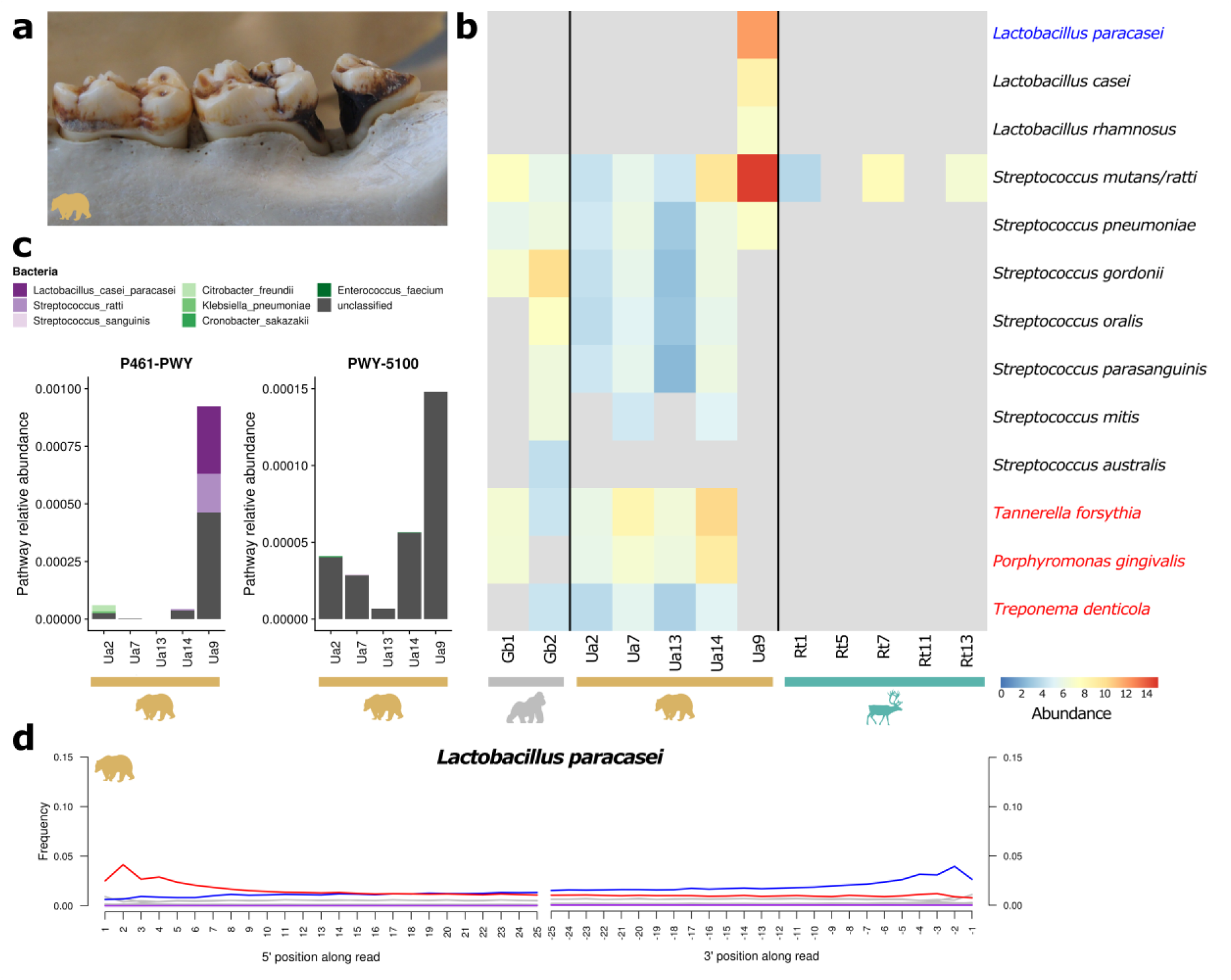
Identification of oral pathogens in a specimen with evidence of oral disease. **a)** Sampling site from a caries lesion from the bear specimen Ua9. **b)** Kraken2 CLR normalised abundance of cariogenic bacteria (*Streptococcus* and *Lactobacillus* species) and periodontal pathogens (*Treponema denticola, Porphyromonas gingivalis* and *Tannerella forsythia*, highlighted in red text). Taxa that were not detected in a sample are coloured grey. **c)** Relative abundance of MetaCyc metabolic pathways involved in sugar fermentation to acids in brown bear samples (P461-PWY: hexitol fermentation to lactate, formate, ethanol and acetate; PWY-5100: pyruvate fermentation to acetate and lactate). Relative contribution of bacterial species to each pathway in each sample is shown where known. Note the differences in y-axis scale for each pathway. **d)** Deamination plot of Ua9 reads mapping to the metagenome-assembled genome *L. paracasei* (highlighted in blue in **(a)**) showing the frequency of C-to-T (red), G-to-A (blue) and all other (grey) substitutions along the 5’ and 3’ ends of the reads. The drop in C-to-T and G-to-A substitution frequencies at position 1 is explained by reduced ligation efficiency to barcoded adapters with terminal C or G bases^40^ (Supplementary Tables 9-10).

By performing shotgun sequencing, we were also able to gain first insights into the functional potential of the microbial communities captured in dental calculus. We used the HumanN2 pipeline^34^ to assign KEGG orthologues^35^ to our filtered microbial reads, and cluster these functions into MetaCyc metabolic pathways^36^. In contrast to the taxonomic results, we observed no clear functional differences between host species (Supplementary Figure 8 and Supplementary Table 2). Instead, we identified core functional pathways shared across individuals and host species that were generally involved in essential metabolic processes, such as energy production and biomolecule synthesis (Supplementary Figure 9), functions common to most living organisms^37^. Conserved functional properties despite differing community compositions is a common phenomenon in ecosystems, including the human microbiome^37,38^. However, this inference could also be driven by limitations of the functional reference databases, which are mostly centred on global mechanisms^39^ and do not allow for more fine-grained functional characterisation. We did identify pathways that have been shown to be enriched in human oral sites, such as those involved in biosynthesis of components of lipopolysaccharide (e.g. ADP-D-glycero-β-D-manno-heptose and lipid A)^37^. Among the basic functions that can be expected from an oral microbial community, we identified a number of metabolic pathways that were particularly abundant in the carious bear Ua9. They were generally involved in carbohydrate fermentation and acid production (Figure 2c), functions commonly performed by bacteria that colonise the oral cavity and are also associated with the emergence and progression of dental caries^33^. Correspondingly, in Ua9, enzymes encoded by *L. paracasei* and mutans streptococci substantially contributed to one of these pathways (P461-PWY, Figure 2c).

The presence of the cariogenic bacterium *L. paracasei* in Ua9 was further confirmed through *de novo* assembly of a high-quality metagenome-assembled genome (MAG) (Supplementary Table 4). The presence of typical DNA damage patterns adds support that the MAG is likely endogenous (Figure 2d). We additionally recovered five other high-quality MAGs from two bear and two reindeer specimens, which had typical DNA damage patterns and were taxonomically classified as strains related to potentially oral bacteria (*Streptococcus* and *Haemophilus*) (Supplementary Table 4). One *Streptococcus* MAG assembled from Ua9 was identified as closely related to *S. ratti*, which likely reflects the high abundance of mutans streptococci identified by Kraken (Figure 2b). Two other high-quality MAGs were recovered from one bear specimen but could not be assigned known taxonomy and thus possibly represent novel bacteria specific to the bear oral cavity.

### Antimicrobial resistance (AMR) genes are present in wild animal microbiomes

The presence of bacteria carrying AMR genes has been documented in the human oral microbiome^10,41^. We therefore investigated whether oral microbial communities of wild animals contain AMR genes and whether their abundance differs across host species. To this end, we investigated both global AMR potential in the entire microbial community and the diversity of AMR genes chromosomally encoded by an oral bacterial pathogen with reported multi-drug resistance, *Acinetobacter baumannii*^42^. To survey global AMR potential, we blasted the contamination-filtered reads against the Comprehensive Antibiotic Resistance Database (CARD)^43^ and assigned the top match for each read to its respective gene family under the Antibiotic Resistance Ontology (ARO) (Figure 3a). To identify AMR genes encoded by *A. baumannii*, we extracted reads aligned by MALT^44^ to the species using MEGAN^45^. We used the EAGER pipeline^46^ to identify reads that mapped to the plasmid-free chromosome and subsequently processed these reads in CARD, as above (Figure 3b). We also confirmed that the identified *A. baumannii* chromosomal reads exhibited characteristic deamination patterns consistent with post-mortem DNA damage (Figure 3c). Both globally and in the *A. baumannii-*specific analysis, gorillas and bears tended to have greater AMR potential compared to reindeer, in terms of both relative abundance (proportion of reads) and diversity (number) of AMR gene families identified. Similar trends were observed in the global dataset when a more conservative, marker-based method of AMR gene family identification was used (Supplementary Figure 10). With our limited sample numbers, we observed no obvious differences in AMR potential in samples collected before and after the start of industrial-scale antibiotics production in the 1940s^47^. However, our analyses are in line with previous studies on historical human and permafrost microbiomes that demonstrate that many of the underlying molecular mechanisms conferring resistance to (modern) antibiotics have existed in the environment long before mass antibiotics production^10,48^. For example, we identified the presence of PER beta-lactamases in *A. baumannii* reads in both Ua7 (dating from 1922) and Ua9 (dating from 1942) (Figure 3b). PER-1 is an extended-spectrum beta-lactamase conferring resistance to third generation cephalosporins and was first observed in *Acinetobacter* species in the late 1990s^49,50^. This investigation opens doors for the use of dental calculus as a tool to study AMR evolution through time in wild animals from diverse geographic locations, and for determining the potential of wildlife to serve as reservoirs for clinically-relevant AMR factors.

**Figure 3.**
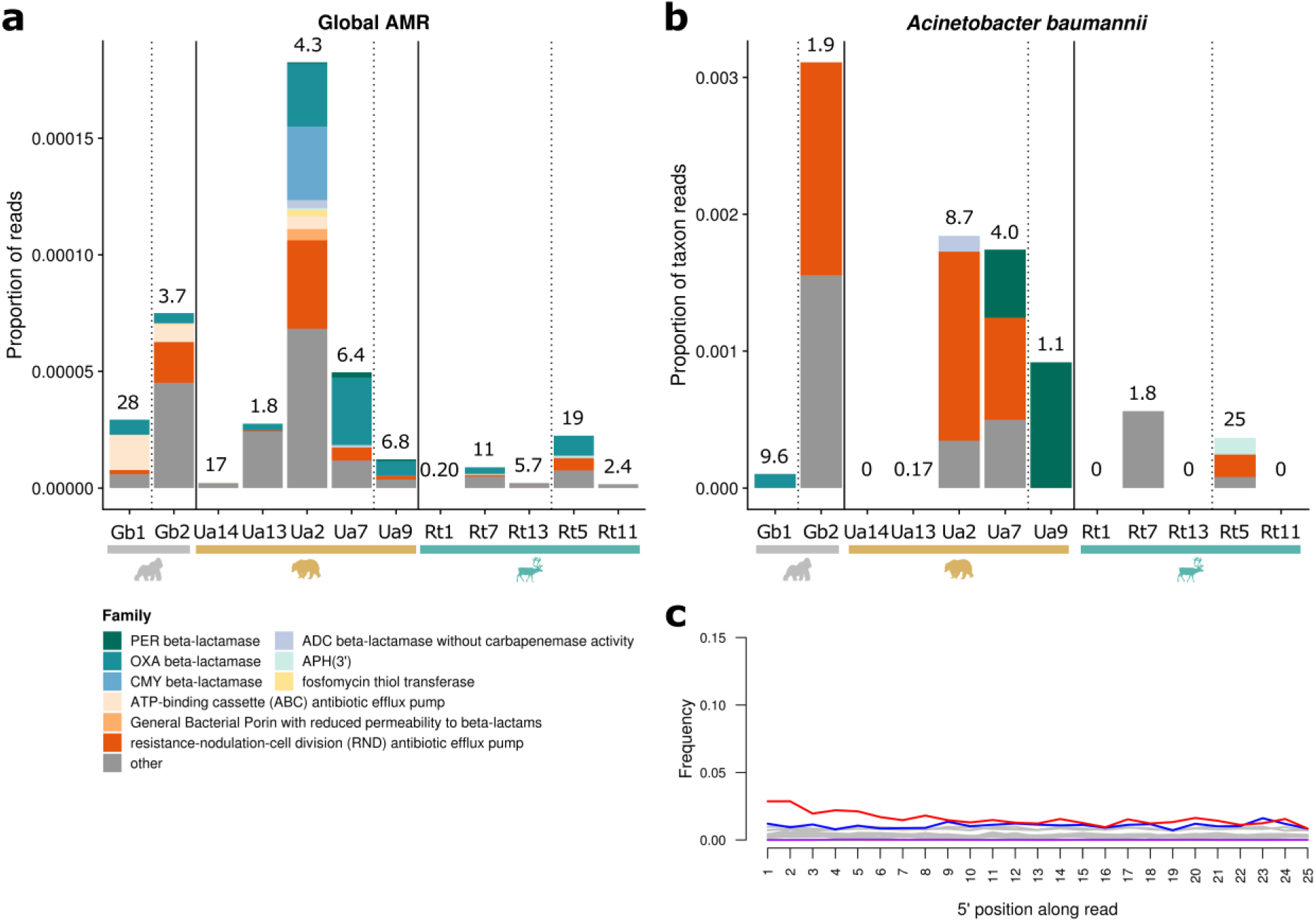
Antimicrobial resistance (AMR) genes can be recovered from historical dental calculus from specimens collected prior to the advent of industrial-scale antibiotics production in the 1940s. **a)** Proportion of global AMR genes identified in all contamination-filtered reads in each sample. **b)** Proportion of *Acinetobacter baumannii* chromosomally-encoded AMR genes identified in each sample. Samples are grouped by host species and ordered by year with pre-1940 samples separated from post-1940 samples by dashed vertical lines. The top nine most abundant AMR gene families are shown, with the remainder grouped into ‘other’. Number of contamination-filtered reads (in millions) is shown above the bars for each sample in (a), and number of reads (in thousands) mapping to the chromosome of *A. baumannii* is shown in (b). **c)** *A. baumannii* is an endogenous member of the mammalian oral microbiome, as evidenced by the presence of typical DNA post-mortem damage patterns exemplified in reindeer Rt5. Frequency of C-to-T (red), G-to-A (blue) and all other (grey) substitutions are shown along the 5’ end of the reads.

### Dental calculus as source of dietary information

To explore the potential of dental calculus to provide insights into the dietary composition of each host species, we taxonomically profiled all eukaryotic reads (excluding those mapping to host and human during data preprocessing) from our samples using MALT^44^ and MEGAN^45^ (Figure 4a). We observed a number of likely spurious mappings and/or contamination, such as to the bovine genus *Bos*, which showed mapping of reads from all three host species and the negative controls (Figure 4b). In bears, only few plant-based dietary components were identified and we found no clear patterns for mammalian or invertebrate putative dietary items, as similar taxa were also present in some reindeer and/or gorilla samples, which are not expected to consume mammals. We were able to infer population-specific dietary characteristics in gorilla and reindeer samples, although in many cases the taxa identified in our analyses are likely close relatives to the consumed species, which are not well represented in the reference databases. For example, among reads that mapped to the Poaceae family in the mountain gorilla sample, approximately 28% mapped to *Phyllostachys*, a genus of Asian giant timber bamboo (Figure 4b). Mountain gorillas are known to consume a related *Arundinaria alpina* bamboo^51^, the genome of which is not currently available. We also identified *Galium* vines in the mountain gorilla (Figure 4b), consistent with the known dietary preferences of Virunga mountain gorillas for which these plants are among the six most frequently consumed taxa^51,52^. *Salicaceae* plants, e.g. *Salix* (willows) were identified in all reindeer (Figure 4b), consistent with the known browsing behaviour of these animals^53^. A number of Arctic plants were identified in the Svalbard reindeer (Rt1 and Rt7) that are known or likely components of the high Arctic reindeer diet, including *Saxifraga* spp. and *Oxyria* spp^54^ (Figure 4b). Furthermore, reads assigned to rumen ciliates from the Ophryoscolecidae family (*Entodinium caudatum* and *Epidinium ecaudatum*) were identified in one of the reindeer samples (Rt11). These protozoa are important facilitators of digestive processes in ruminants^55^. Similarly, in the microbial analyses, several microbial taxa specific to the reindeer were identified as *Methanobrevibacter* species associated with the rumen of domesticated bovines and ovines^56^. Ruminants regurgitate large amounts of rumen material into the oral cavity when chewing cud, and a study in sheep has found that oral swabs contain a proportion of rumen-associated microbes^57^. It is therefore plausible that dental calculus of ruminants also captures a subset of the rumen microbiome.

**Figure 4.**
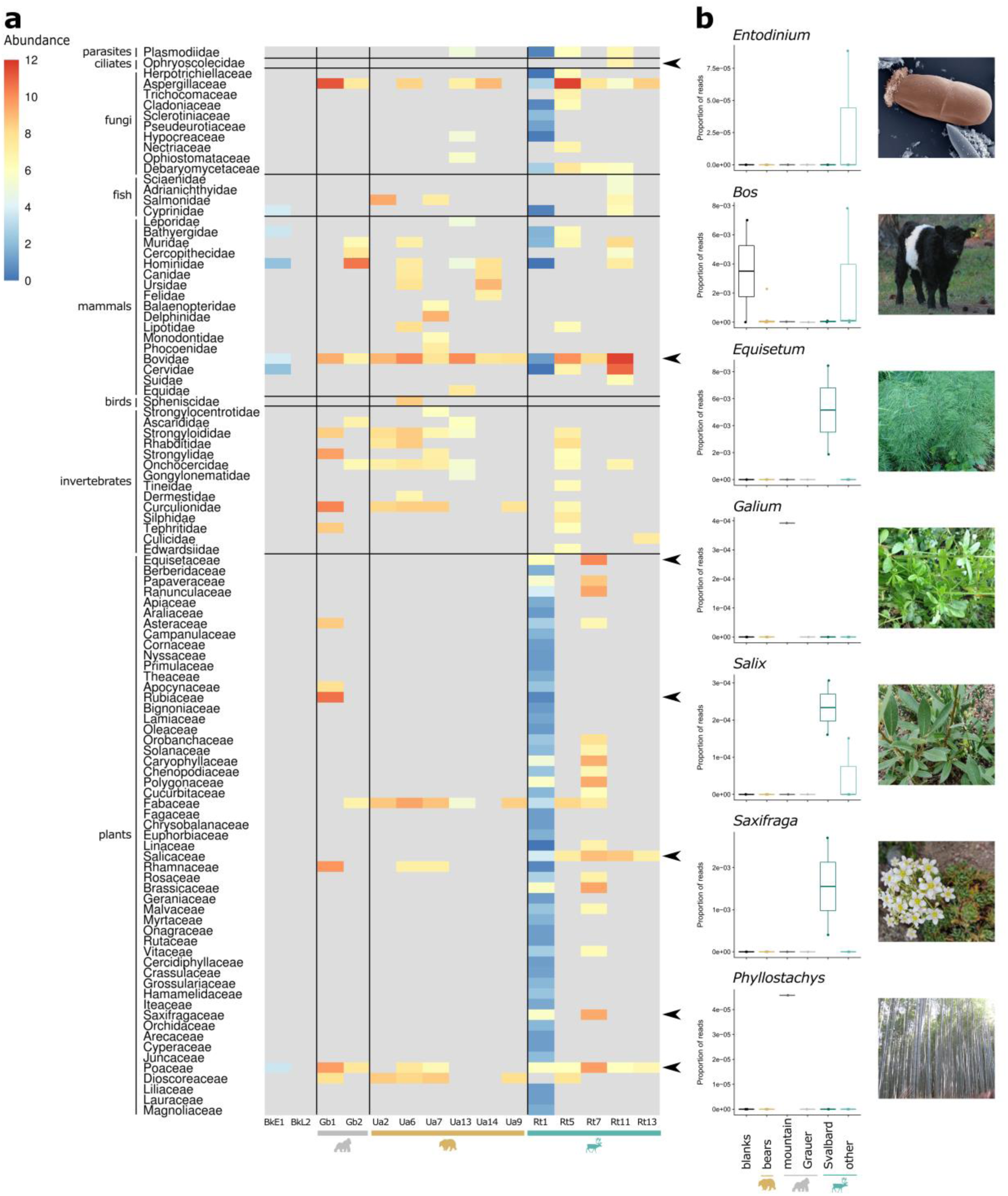
Host diet can be inferred from dental calculus. **a)** MALT/MEGAN CLR normalised abundance of eukaryotic reads at the family taxonomic level. Taxa that were not detected in a sample are coloured grey. Broad groups of eukaryotes are designated by horizontal black lines. Black arrows indicate selected families for which genus-specific relative abundances were plotted in (b). **b)** Proportion of reads mapping to specific eukaryotic genera in different host species, including blank controls (visualised as Tukey boxplots). Gorilla samples are divided into the two subspecies (mountain and Grauer’s gorillas) and reindeer are divided into the Svalbard ecotype and “other” ecotypes (mountain and forest) to illustrate population-specific differences in dietary components. The Bovidae genus *Bos* is included as an example of spurious mappings, due to the presence of reads in samples from multiple host species and blank controls. Image credit: *Entodinium caudatum* photo by Sharon Franklin and colourisation by Stephen Ausmus (https://www.ars.usda.gov/oc/images/photos/feb06/d383-2/), other images by Katerina Guschanski and Jaelle Brealey.

### Recovery of host genomic profiles

In addition to microbial remains, dental calculus also entraps host cells and thus can serve as a source for host DNA^12,58^. We therefore identified host DNA preserved in dental calculus samples (including bear Ua6 excluded from the microbial analyses) by mapping the reads to reference genomes of the host species’ closest phylogenetic relatives. For mitochondria, 2.6-99.7% (median 91.2%) of the genome was covered by at least one read in each study sample with the coverage depth ranging from 0.02 to 174x (Supplementary Table 5). Indeed, the high abundance of host reads in some samples allowed us to reconstruct complete mitochondrial genomes from five specimens (Supplementary Table 5). For nuclear genomes, 0.004-21.3% (median 0.346%) were covered by at least one read, with maximum genome-wide coverage of 0.3x. We compared the recovered genomic profiles to published genomes from the same species. Mitochondrial haplotypes could be placed within species-specific mitochondrial networks (Figure 5a and Supplementary Figures 11-12, note that only a single reference genome is currently available for reindeer, which limited our inferences for this species). In bears, the mitochondrial haplotype network reflected known differences in colonization history of Scandinavia from west and east^59^: one of the two Swedish bear samples (Ua14 from central Sweden) was placed together with central European bears and the other (Ua9 from northern Sweden) with Russian/Alaskan individuals (Figure 5a). Projection of low-coverage host nuclear genomes onto the PCA space pre-calculated from high-quality published genomes clearly assigned all study samples to their correct species (reindeer), subspecies (gorilla), and even to broader geographic populations of origin (brown bear) (Figure 5b-d).

**Figure 5.**
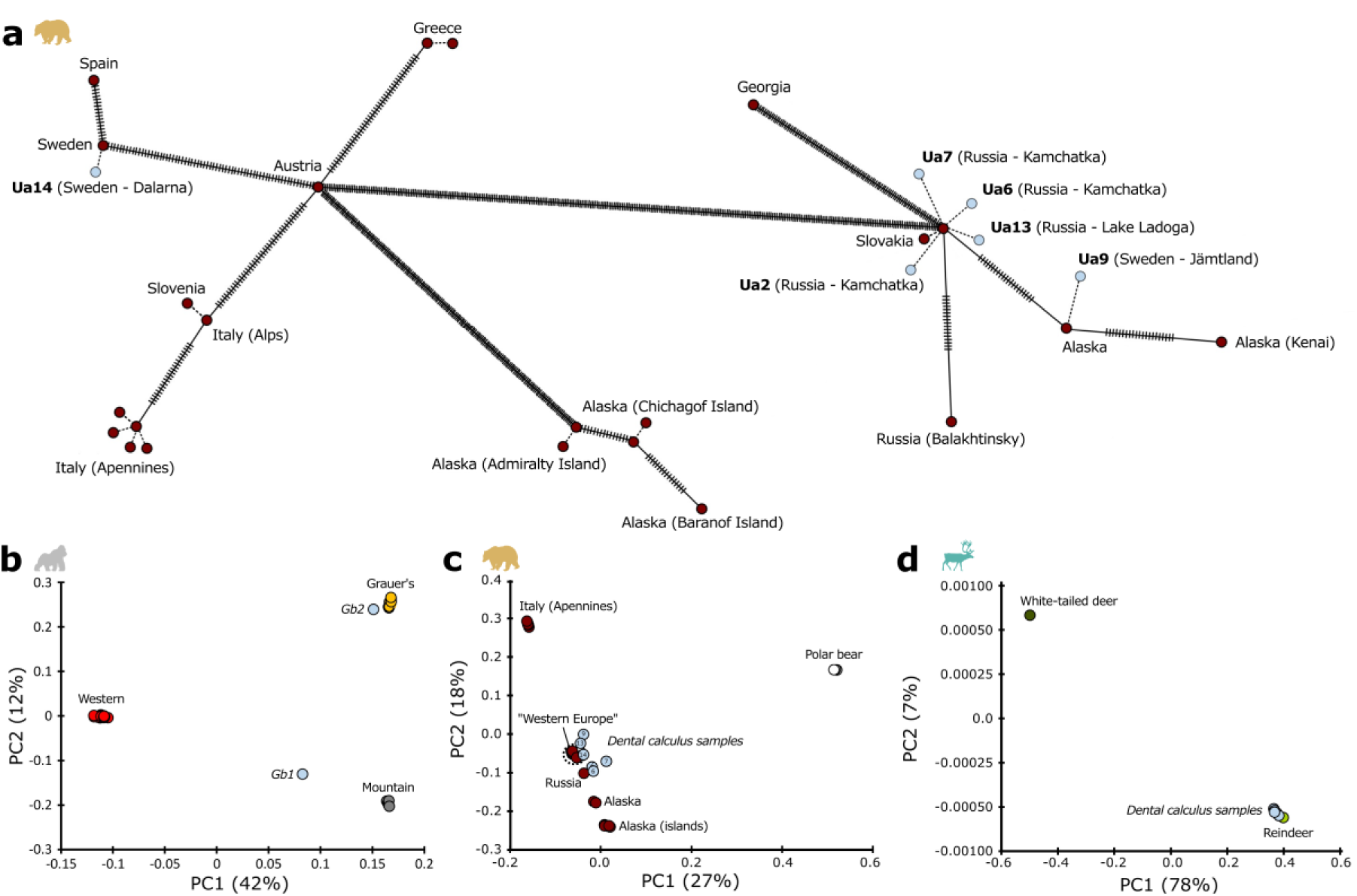
Host population genetic structure can be reconstructed from dental calculus. **a)** mtDNA haplotype network for brown bears. Each circle represents a sample, with ticks on the connecting lines showing the number of base pair substitutions between the haplotypes. Dotted lines represent identical haplotypes or in the case of dental calculus samples (shown in light blue), the predicted most closely related haplotype. Sample labels include country and locality of specimen collection. **b-d)** PCA of modern high-coverage genomes and low-coverage dental calculus samples projected onto the modern genomes for gorilla (b), brown bear (c) and reindeer (d). Samples cluster together with their respective species. Dental calculus gorilla samples cluster most closely to their subspecies of origin. Dental calculus brown bear samples from western Europe and Russia cluster with the modern genomes from the “Western Europe” clade (consisting of Spain, Greece, Slovenia, Italy (Alps), Sweden and Slovakia) and the Russian clade, respectively. Dental calculus reindeer samples cluster with the single available reindeer reference genome.

### Dealing with current limitations: Contamination and database biases

Dental calculus has been little explored outside of human research, yet it can be a treasure trove of information about evolutionary and ecological processes of the host and its oral microbiome. However, research on microbiomes from the past, including from dental calculus, is hindered by a number of challenges. We have established rigorous methodology for overcoming the problem of contamination, which affects all historical genomic and metagenomic studies^18^. Through a combination of laboratory and computational procedures, we reduced the proportion of contaminants and systematically increased the proportion of the bacterial communities attributed to the oral microbiome (Supplementary Figure 1). However, the SourceTracker results indicated that some proportion of human skin taxa remained in our samples. Several bacterial species colonise multiple niches within the host^37,38^, which can obscure distinction of a genuine signal from a likely contaminant. For example, *S. mitis*, *Staphylococcus epidermidis* and *Corynebacterium matruchotii* are found in the human mouth, nostrils and skin^38^. This limitation is not specific to our dataset, but poses greater problems for studies based on historical samples that are expected to be subject to contamination. The other common limitation faced by our and many other studies is the reliance on microbial reference databases. These databases are heavily biased towards microbial species with medical or agricultural significance^60^, restricting read-based analyses of metagenomics data from non-human hosts. A large proportion of microbial taxa identified in our samples remained unassigned to any source microbiome by SourceTracker. Although some of these taxa may be members of other microbial communities not included as a source in our analysis, we expect that by studying a novel environment (the non-human oral microbiome) we will encounter unique microbial taxa. In the absence of a dedicated reference database from the study species, an alternative approach is *de novo* MAG assembly^60,61^. Given the fragmented and damaged nature of ancient DNA, this technique poses great challenges for historical microbiome studies. However, our study demonstrates that with deeper sequencing MAG recovery may be able to complement read-based analyses of historical microbiome samples. This is particularly important in non-model species, where reference database bias is a problem.

## Discussion

With the development of high-throughput sequencing techniques and methodological advances in metagenomic analyses of ancient samples, the time is ripe to venture into the study of the diversity of environmental and host-associated microbial communities from the past. The temporal perspective provided by historical and ancient samples allows us to study many fundamental evolutionary processes, including those with direct relevance to human and ecosystem health. Our study describes a rigorous roadmap for the analysis of historical microbiomes and illuminates a magnitude of biological questions that can benefit from the study of dental calculus remains. We demonstrate that a single sample source can be used to link the host microbial community to host genetics, diet and even disease. While larger sample sizes are needed to substantiate the biological inferences of our preliminary findings, our study establishes dental calculus as a tool for the exploration of a broad variety of research topics. Questions of interest include the evolution of host-associated microbiome through multi-generational changes in the host environment, such as climate change, alterations in host population demography and genetic diversity, the invasion of new habitats, or changes in competitive regimes. Temporal sampling of dental calculus also provides insights into oral disease emergence and the progression of antimicrobial resistance in host-associated microbiomes. These processes can be of interest to both evolutionary biologists and the public health sector, since wild animal populations can act as sources and reservoirs for emerging zoonotic pathogens^62,63^ and contribute to the spread of antimicrobial resistance^64,65^. In addition, our dietary results indicate that dental calculus can be used to infer population-specific dietary characterises, particularly if complemented with microfossil analysis from the same material and stable isotope analysis of teeth or bones^66–68^, which can be extended to extinct species (e.g.^69^). We envision that in the future, our integrative approach will be applied to many different questions in ecology and evolution.

## Materials and methods

### Sample collection

Dental calculus was collected from two eastern gorilla (*Gorilla beringei*) specimens from the Royal Museum for Central Africa (Brussels, Belgium), as well as five reindeer (*Rangifer tarandus*) and six brown bear (*Ursus arctos*) specimens from the Swedish Natural History Museum (Stockholm, Sweden). Skulls were macroscopically examined for dental calculus deposits and evidence of oral diseases. Calculus was removed from the surfaces of the teeth with disposable sterile scalpel blades and deposited in sterile microcentrifuge tubes.

### Sample processing and DNA extraction

All laboratory protocols were performed in a dedicated ancient DNA laboratory following stringent procedures to minimise contamination^18^. Initially, calculus samples were processed without surface decontamination (Supplementary Table 1). We then tested the effect of surface contamination on the two gorilla samples using three treatments: exposing the calculus to UV light (245 nm) for 10 min, washing the calculus in 500 µl of 0.5 ethylenediaminetetraacetate (EDTA) for 30 seconds, and no surface decontamination as a control. Based on real-time PCR of libraries prepared from these samples (see below), we determined that neither decontamination treatment substantially reduced library quantity and thus continued with a surface decontamination procedure consisting of the UV light exposure followed by the EDTA wash for all subsequent calculus samples (Supplementary Table 1). The pellet was taken forward for DNA extraction. Sample weight ranged from < 5 mg to 20 mg. DNA extractions were performed following a silica-based method^70^. Briefly, samples were incubated overnight at 37^°^C in extraction buffer (0.45M EDTA, 0.25 mg/ml Proteinase K). The DNA from the supernatant was combined with binding buffer (3M sodium acetate, 5M guanidine-hydrochloride, 40% (v/v) isopropanol, 0.05% (v/v) Tween-20) and processed through the spin columns from High Pure Viral Nucleic Acid Large Volume kits (Roche, Switzerland). Purified DNA was eluted in either 45 µl of EB buffer (10 mM tris-hydrochloride (pH 8.0) (Qiagen, The Netherlands) or 45 µl of TE buffer (10 mM tris-hydrochloride (pH 8.0), 1 mM EDTA), both supplemented with 0.05% (v/v) Tween-20 (Supplementary Table 1). Ten blanks were included during five extraction batches and carried through library preparation.

### Library preparation and sequencing

Double-stranded Illumina libraries were prepared following ^71^ and we included a double-barcoding double-indexing strategy to guard against index hopping and retain absolute certainty about sample of origin^40,72^. Briefly, blunt-end repair reactions were performed using 20 µl of each extract and purified using MinElute columns with elutions in 22 µl of EB buffer (Qiagen, The Netherlands). Adapters containing inline 7 bp barcodes (Supplementary Tables 1 and 6) were ligated to both ends of the blunt-ended DNA, which was subsequently purified with MinElute columns and eluted in 22 µl of EB buffer. After the adapter fill-in reaction, *Bst* 2.0 polymerase (New England BioLabs, USA) was inactivated with a 15 min incubation at 80°C. Seven library blanks were included in four library preparation batches. The adapter-ligated libraries were quantified using a real-time PCR assay with preHyb primers^40^ (Supplementary Table 6) and the estimated fragment number was used to approximate the number of indexing PCR cycles needed for sequencing. All extraction and library blanks were consistently lower in DNA content than samples, as measured by real-time PCR, thus one extraction blank and one library blank were randomly selected for subsequent indexing and sequencing. Libraries were double-indexed with unique P5 and P7 indices so that each sample had a unique barcode-index combination (Supplementary Table 1). Indexing PCR reactions were performed with 18 µl of adapter-ligated library in 50 µl reactions, with 1 µl PfuTurbo Cx hotstart polymerase (2.5 U/µl, Agilent Technologies, USA), 5 µl 10X PfuTurbo Cx reaction buffer, 0.5 µl dNTP mix (25 mM) and 1 µl of each indexing primer (10 µM). After an initial incubation for 2 min at 95°C, 12 cycles of 30 sec at 95°C, 30 sec at 59°C and 1 min at 72°C were performed, followed by a final step of 10 min at 72°C. Reactions were purified with MinElute columns and eluted in 10 µl of EBT buffer. The indexed libraries were quantified using a real-time PCR assay with i7 and i5 indexing primers^40^ (Supplementary Table 6) and library DNA fragment length distribution was determined by the 2200 TapeStation system. The mean fragment length after library preparation and excluding the 148 bp adapter sequences was 70 bp, similar to what has been observed in previous historical sequencing libraries^12,73^. Five microliters of each sample library were pooled along with a randomly selected extraction blank and library blank. Size selection was performed on the pooled library with AMPure XP beads (Beckman Coulter, USA), selecting for fragments approximately 100-500 bp in length, and the purified library eluted in 36 µl of EB buffer. The final pooled library was quantified using a Qubit High Sensitivity fluorometer and on the 2200 TapeStation system. The pooled library was first sequenced by SciLifeLab Uppsala on 2 lanes of the Illumina HiSeq 2500 using paired-end 125 bp read length v4 chemistry, followed by an additional 2 lanes on the Illumina HiSeq 2500 in rapid mode using paired-end 100 bp read length v2 chemistry.

### Data processing

Sequencing data were demultiplexed and assigned to each sample with an in-house python script based on the unique combination of barcodes and indices. Overlapping paired-end reads were merged and adapters and low quality terminal bases (phred scores ≤ 30) were removed with AdapterRemoval v2.2.2^74^. Barcode sequences were removed from the 5’ and 3’ ends of merged reads with an in-house python script. Forward reads from the unmerged read pairs (i.e. pairs that did not contain overlapping regions of at least 11 bp between the forward and reverse reads) were also retained for analyses. The 5’ barcode was removed with an in-house python script and the 3’ barcode with any remaining adapter sequence removed with AdapterRemoval. Reads from the two lanes within the same sequencing run were concatenated into a single file per sample. Merged reads from the two separate runs were also concatenated into a single file per sample. Reads with a length < 30 bp were filtered out with AdapterRemoval and reads with mean base quality < 30 were filtered out with PrinSeq-Lite v0.20.4^75^. Duplicate reads were removed by randomly keeping one read among those reads having an identical sequence. The Illumina sequencing control phage PhiX was spiked into our sequencing runs and has been reported to have been erroneously integrated into many microbial reference genomes^76,77^. Reads were therefore mapped to PhiX (accession: GCA_000819615.1) with bwa mem v0.7.17^78,79^ and the unmapped reads retained with SAMTools v1.9^80^ and BEDTools v2.21.0^81^. To remove reads originating from the host organism and from human contamination, we mapped all reads in a sample to a combined reference consisting of the human genome^82^ (RefSeq accession: GCF_000001405.38) and the respective host genome (GCF_000151905.2 (*Gorilla gorilla gorilla*)^83^, GCF_003584765.1 (*U. arctos horribilis*)^84^ and GCA_004026565.1 (*Rangifer tarandus*)) with bwa mem. The unmapped reads were retained with SAMTools for downstream microbial taxonomic analyses.

### Taxonomic assignment

Merged and unmerged unmapped reads were assigned taxonomy using the *k*-mer based classifier Kraken2 v2.0.7^19^ with the standard Kraken2 database (all archaea, bacteria, viruses and the human genome in RefSeq; built 2019-03-01). We used Kraken-biom (https://github.com/smdabdoub/kraken-biom) to extract the summarised number of reads assigned at the genus and species levels. These assignments were taken into R (https://www.R-project.org/) for further processing and analysis.

We also taxonomically binned the contamination-filtered reads (see below) against a wider database by alignment with MALT v0.4.0^44^ (parameters as per ^12^: semi-global alignment, 85% minimum identity threshold, 0.01% minimum support threshold and a top percent value of 1.0) against an index built from the entire NCBI nucleotide database (ftp://ftp.ncbi.nlm.nih.gov/blast/db/FASTA/nt.gz, modified 31-05-2018) using a step size of 4 to reduce index size. The results were viewed in MEGAN Community Edition v6.10.5^45^. Due to relatively low specificity leading to spurious alignments that required manual curation, we did not use the MALT/MEGAN results for microbiome analyses, but took them forward for specific bacterial alignments as well as for eukaryotic taxonomic assignments (see below).

### Identifying contamination

We used several complementary approaches to identify and remove contaminating bacterial taxa from the Kraken taxonomy assignments.

#### SourceTracker analysis

Potential contribution of source microbiomes to samples was estimated with SourceTracker v1.0^22^ in R, using the Kraken2 genus- and species-level assignments. Source sequencing reads were processed through the same pipeline as sample reads, and included soil^85^, human skin^86^, human gut^37,38^, human supragingival plaque^37,38^, human medieval dental calculus^12^ and laboratory reagent^20^ microbiomes (Supplementary Table 7).

#### Fragment length

Given our library fragment length distribution peak at 70 bp, consistent with historical degraded DNA, and our paired-end sequencing approach of 100 bp, we expected that the majority of reads stemming from authentic historical microorganisms would be successfully merged by AdapterRemoval. In contrast, modern contaminating taxa with longer fragment lengths should be predominantly found among the unmerged reads. We therefore compared the raw read counts of each taxon between the merged and unmerged reads on a per sample basis (Supplementary Figure 2). The difference in read number between the unmerged and merged reads for each taxon in each sample was calculated and the distribution of all differences > 0 (i.e. more reads observed in the unmerged reads) was investigated (Supplementary Figure 2). Taxa identified as outliers (defined as a difference > 1.5 interquartile ranges above the third quartile) in at least one sample were filtered out as putative contaminants. While it is possible that certain microbial structural features, e.g. the thick cell wall in Gram-positive bacteria, might bias preservation towards specific taxa and thereby affect our fragment length analysis^18^, a recent study of microbial DNA perseveration in human dental calculus found no association between cellular structures and microbial DNA fragmentation^12^. Nonetheless, we confirmed that the majority of taxa in our putative contaminant list were non-host-associated environmental microorganisms through a literature search.

#### Statistical identification of contaminants

Contamination derived from environmental sources (including the laboratory) is expected to be present in samples at approximately similar absolute amounts and will therefore be disproportionally more abundant in samples with low numbers of total sequenced reads^20,87^. This concept has been implemented in the R package decontam^21^, which was used to identify and remove all taxa that showed an inverse relationship between taxon abundance and total number of sequences included in the sequencing pool per sample, as estimated by real-time PCR.

#### Presence in blanks

SourceTracker analysis demonstrated that the two blank samples contained taxa associated with soil and human skin microbiomes (Supplementary Figure 1). However, low levels of sample cross-contamination are common during laboratory processing. Conservatively filtering out all taxa observed in the blanks might remove genuine signals. We therefore screened all taxa that were present in the blanks against the Human Oral Microbiome Database (HOMD)^88^. Taxa that were not present in HOMD were classified as contaminants and removed from further analysis.

#### Removal of contaminants from reads

For all downstream analyses, reads mapping to all species-level taxa (bacteria, archaea and viruses) identified as putative contaminants were removed from the fastq files. To this end, one reference genome for each taxon was downloaded from GenBank (Supplementary Table 8; accessed 2019-04-11) and reads were mapped to the reference genomes with bwa mem v0.7.17, relaxing the mismatch parameter (-B) to 3 in order to map reads from closely related strains. Only unmapped reads were passed onto further analyses.

### Microbial analyses

#### Genome size normalisation

Taxa with larger genomes will generally contribute more sequencing reads to a library, biasing read-based abundance estimates^89^. To account for this bias, the number of reads per taxon within a sample was normalised by dividing by the estimated average genome size of each respective taxon. Estimated average prokaryotic and viral genome sizes were calculated using publicly available genome sizes from the RefSeq database (accessed 2019-02-15)^90^. In cases where no genome size data were available for a given species, the average genome size of taxa in that genus was used.

#### Abundance filtering

Taxa present at < 0.03% relative abundance (normalised read count divided by sum of normalised read counts in a sample) were removed, as filtering of low-abundance species reduces false-positive taxonomic assignments^91^. The filtering threshold was selected by testing a series of thresholds commonly applied in metagenomics studies, ranging from 0.01–0.1%^92^(Supplementary Figure 13). From this analysis, we identified the threshold (0.03%) that yielded a microbial community with a complexity which was most similar to what has been observed in other dental calculus^12^ and oral microbiome studies^93–96^ (i.e. approximately 100-300 taxa).

#### Abundance normalization

In high-throughput microbiome sequencing datasets, the total number of reads obtained is an arbitrary value set by the sequencing instrument and absolute abundance of each taxon is unknown^97^. Therefore, to account for the compositional nature of the data, we applied the centred log-ratio (CLR) transformation^98^. Because log transformation is only possible for positive values, we dealt with 0 count values by adding a pseudo-count to the normalised read count for every taxon in every sample. Due to the genome size normalisation, we set the pseudo-count to the equivalent of one read divided by the average genome size for all taxa.

#### Statistical analyses

We used the R package vegan^99^ for diversity estimates. The Shannon index^100^ was used to estimate alpha diversity, via the diversity function on the raw read count data (i.e. prior to genome size normalisation and CLR transformation), because the calculation of the metric required positive integers. Differences between host species were investigated with an AIC-based stepwise regression to determine the best-fit general linear model, with surface decontamination, sequencing depth (number of unmapped reads per sample) and proportion of human reads per sample included as covariates.

Beta diversity (a measure of inter-individual variation) was investigated based on the presence/absence of microbial taxa. Jaccard distances, calculated using the vegdist function, were used for ordination with non-metrical multidimensional scaling (NMDS), and permutational multivariate analysis of variance (PERMANOVA) with the adonis function in vegan. Host species, surface decontamination, number of unmapped reads per sample, and proportion of human reads per sample were included as covariates in the adonis model. To determine whether differences in within-group variation between host species was biasing inferences of a host-specific oral microbiome signature, a distance-based test for homogeneity of multivariate dispersions was performed with the vegan function betadisper. No such differences were detected, adding confidence to the PERMANOVA results. The same set of beta analyses were repeated using Euclidean distances calculated on the CLR normalised abundance data.

To identify taxa which discriminated between host species, we carried out a random forest classification based on presence/absence data using the R package randomForest^101^ with 10000 trees. This approach reports the out-of-bag estimated error (how often an individual was incorrectly assigned to a host species) and the variable importance (mean decrease in accuracy) of each taxon, which reflects the importance of the given taxon in determining the correct host species. For this analysis, the gorilla samples were excluded due to low sample size (n=2). We also investigated a subset of taxa unique to each host species and shared by > 50% of samples from this species to determine whether they were likely oral taxa. As the two gorilla samples did not share any taxa, we investigated all of the taxa unique to each of the two samples. We then compared these sets of taxa to the HOMD^88^ and classified those present as ‘oral’. Taxa not present in the HOMD were manually classified as ‘oral’ (based on the presence in the oral microbiome of non-human mammals), ‘host-associated’ (present in non-oral mammalian microbiomes) or ‘not host-associated’ through a literature search.

### Functional analysis

#### Classification

The functional genic content of the microbial community in dental calculus was characterised by running the contamination-filtered reads through the HUMAnN2 pipeline^34^, which identifies species-specific genes with the taxonomic profiler MetaPhlAn2^102^ and a built-in microbial pangenome database representing all known nonredundant protein-coding potential for each species identified by MetaPhlAn2, and more general functional characterisation by alignment with DIAMOND^103^ against the UniRef90^104^ database. The mappings are weighted by quality and sequence length to estimate species-specific and total community gene family abundance. Metabolic pathways are also reconstructed based on genes annotated to metabolic enzymes in MetaCyc^36^, and the pathway abundance and coverage are reported.

CLR normalisation, NMDS ordination and PERMANOVA were carried out for the pathway abundances as for the microbial analyses outlined above. Core pathways were defined as those containing > 50% of the required enzymes in a sample (i.e. per sample pathway coverage > 0.5) for the total community (i.e. not stratified by microbial species). Relative (proportional) abundance for specific pathways stratified by microbial species was also calculated by HUMAnN2 for each sample.

### Antimicrobial resistance profiles

#### Global

Contamination-filtered reads were aligned to the Comprehensive Antibiotic Resistance Database (CARD) v3.0.1 (modified 2019-02-19)^43^, a curated collection of resistance determinant sequences, with blast v2.7.1+^105,106^ using default parameters. The Antibiotic Resistance Ontology (ARO) accession number associated with each CARD sequence was used to obtain the AMR gene family of each sequence. Where reads matched multiple sequences in the CARD, the best hit was identified based on highest bit score. Where multiple hits had the same bit score, we compared the ARO terms and if all hits shared the same ARO information, we randomly chose one hit to carry forward. When ARO information was not identical, we manually identified a common higher level term of the hits: for example, if all hits were to different beta-lactamases, we referred to the gene family as “unclassified beta-lactamase”. We then calculated the number of reads per ARO accession per sample and normalised it by sample sequencing depth (number of reads post-contamination filtering). The abundance of ARO gene families for each sample was calculated by summing across the ARO accession abundances associated with each gene family.

#### Targeted

All complete genomes for *A. baumannii* (n=133) available at NCBI Genome (accessed 2019-03-28) were downloaded and the fasta files concatenated into one file representing the ‘pangenome’. The plasmid sequences were removed to exclude analysis of reads that map to plasmids, since plasmids can be exchanged between bacterial species through horizontal gene transfer, which can lead to erroneous taxonomic identification. Reads assigned by MALT to *A. baumannii* on at least the species level were identified in each sample by MEGAN. These reads were then mapped to the plasmid-free pangenome with bwa mem. All mapped reads were extracted with SAMTools (samtools fastq-F4) and aligned to CARD as per the global AMR analysis outlined above. The abundance of *A. baumannii* ARO gene families for each sample was calculated by summing across ARO accession abundances normalised by number of reads mapping to each bacteria.

#### Marker-based

Contamination-filtered reads were also compared to a set of AMR gene family marker sequences built via ShortBRED^107^ from CARD v3.0.1 (modified 2019-02-19) using UniProt Reference Cluster 90 (UniRef90, modified 2019-05-08)^108^ as the background. Normalised counts (reads per kilobase of reference sequence per million sample reads) for each family in each sample were subsequently analysed as above.

### Metagenome-assembled genome (MAG) recovery

We attempted to recover MAGs from our samples following a similar strategy as described in Zhou *et al.* 2018^109^ and Parks *et al.* 2017^61^. Reads were assembled into contigs with MEGAHIT^110^. Sample depth profiles were generated by mapping reads back to the contigs with BBMap v38.08 (Bushnell B., *BBMap*, https://sourceforge.net/projects/bbmap/). Contigs > 1500 bp in length were grouped into genome bins based on co-abundance and sequence composition probabilities using MetaBAT2^111^, where each bin theoretically represents the genome of a single strain. The quality of each bin was assessed using the lineage-specific workflow in CheckM^112^. High quality bins were classified as those with an estimated > 90% genome completeness and < 5% strain contamination^61^. The taxonomic lineage of each high quality bin, as defined in the Genome Taxonomy Database (GTDB)^113^, was determined with GTDB-Tk (https://github.com/ecogenomics/gtdbtk) using the classify workflow.

### mapDamage profiles

Deamination rates and plots were generated using mapDamage^114^ via the EAGER pipeline^46^. During this investigation, we observed unusual damage patterns in several of our samples – rather than the expected incremental rise in deamination towards read ends, several samples showed a drop in the frequency of C-to-T substitutions from the penultimate base to the terminal base (e.g. Figure 2d [*L. paracasei* from Ua9] compared to Supplementary Figure 3 [*Streptococcus* sp. Z15 from Rt7]). It has been suggested previously that barcodes with a terminal G or C have reduced ligation efficiency to damaged cytosines at 5’ ends of reads^40^. We therefore examined damage profiles for 14 most abundant bacterial taxa in the post-filtering dataset (Supplementary Table 9), only considering cases with more than 10000 reads mapping to a taxon reference genome in a given sample. We found that samples containing barcodes with a terminal G or C were more likely to show the unusual pattern (χ^2^ = 31.11, df = 1, p-value < 0.0001) (Supplementary Table 10).

### Dietary component recovery

Counts of reads assigned by MALT to eukaryotic taxa were exported from MEGAN at the family and genus level and imported into R. Taxa with <10 assigned reads across all samples were excluded from further analysis. At the family level, assigned read counts were normalised across samples by adding a pseudo-count of 0.1 and applying the CLR transformation. The proportion of specific genera per sample was calculated by taking genus level read counts and normalising by sample sequencing depth.

### Host genome recovery

We collected published whole genome sequence data for 1 reindeer (GCA_004026565.1), 1 white-tailed deer (GCA_002102435.1), 47 gorillas^115–117^, 24 brown bears and 3 polar bears^118–121^. Reads were mapped either merged (ancient samples), or paired-end (modern samples) to an outgroup reference genome assembly (white-tailed deer: GCA_002102435.1, human: GCA_000151905.3 and polar bear: GCA_000687225.1) using bwa mem on default settings. Next, we excluded reads with a mapping quality score < 30 and removed duplicate reads with SAMTools v0.1.19. Additionally, for each study host species, we mapped all reads to the mitochondrial references of white-tailed deer (NC_015247.1), polar bear (NC_003428.1) and western gorilla (NC_011120.1), respectively, following the same pipeline.

To investigate host genomic variation, we generated pseudo-haploid sequences for each individual by randomly selecting a single high-quality base call (BaseQuality ≥30, MapQuality ≥30) at each site covered by at least 1 read, excluding sites within repetitive regions (as identified from the repeatmask tracts) and for modern genomes sites with >2 times genome-wide coverage to minimize false bases from spurious mappings. A reference set of high quality polymorphisms was made from all bi-allelic autosomal sites in the modern genomes and we then projected the low-coverage dental calculus samples onto the precalculated PCA space using the lsqproject function in smartpca^122^.

Mitochondrial genome variation was investigated following the same pipeline but calling the majority allele at each covered site. We then used popart^123^ to create haplotype networks for all near-complete mitochondrial genomes (>90% complete). For the incomplete mitochondrial genomes, we calculated pairwise-divergence to each of the modern genomes to obtain the most likely closest related haplotype.

In addition we also obtained *de novo* mitochondrial genomes using MITObim v1.9^124^, a mitochondrial baiting and iterative mapping method, which reconstructs complete mitochondrial genomes from next generation sequencing data. The mitochondrial genomes of the western lowland gorilla (*Gorilla gorilla gorilla,* NC_011120.1), the brown bear (*Ursus arctos,* NC_003427.1) and the reindeer (*Rangifer tarandus*, KM506758.1) were used as bait sequences for the mitochondrial genome assembly of the eastern gorilla, brown bear and reindeer samples, respectively. The merged host reads were used as input files for each sample. All mitochondrial genomes were annotated with the MITOS WebServer^125^ using default parameters.

### Data availability

All sequencing data are archived at the European Nucleotide Archive under the accession number PRJEBxxxxx. Sample metadata is provided in Supplementary Table 1.

## Acknowledgements

We thank Daniela Kalthoff and Emmanuel Gilissen for providing access to the mammalian museum specimens, Peter Niehoff and Riana Minocher for assisting with sampling, and James Fellows Yates, Franziska Aron, Johanna von Seth and Tatiana Feuerborn for laboratory and analysis advice. Sequencing was performed by the SNP&SEQ Technology Platform in Uppsala. The facility is part of the National Genomics Infrastructure (NGI) Sweden and Science for Life Laboratory. The SNP&SEQ Platform is also supported by the Swedish Research Council and the Knut and Alice Wallenberg Foundation. We also acknowledge the National Bioinformatics Infrastructure (NBIS) for providing computational resources to this project. The project was supported by the Formas grant (2016-00835) and the National Projects grant (NP00039) to KG.

