## Supplemental Figures S1-S12 and Supplemental Tables S1, S3-S7, S10 for "A roadmap to mammalian oral microbiome evolution with dental calculus"

### Supplementary Figures

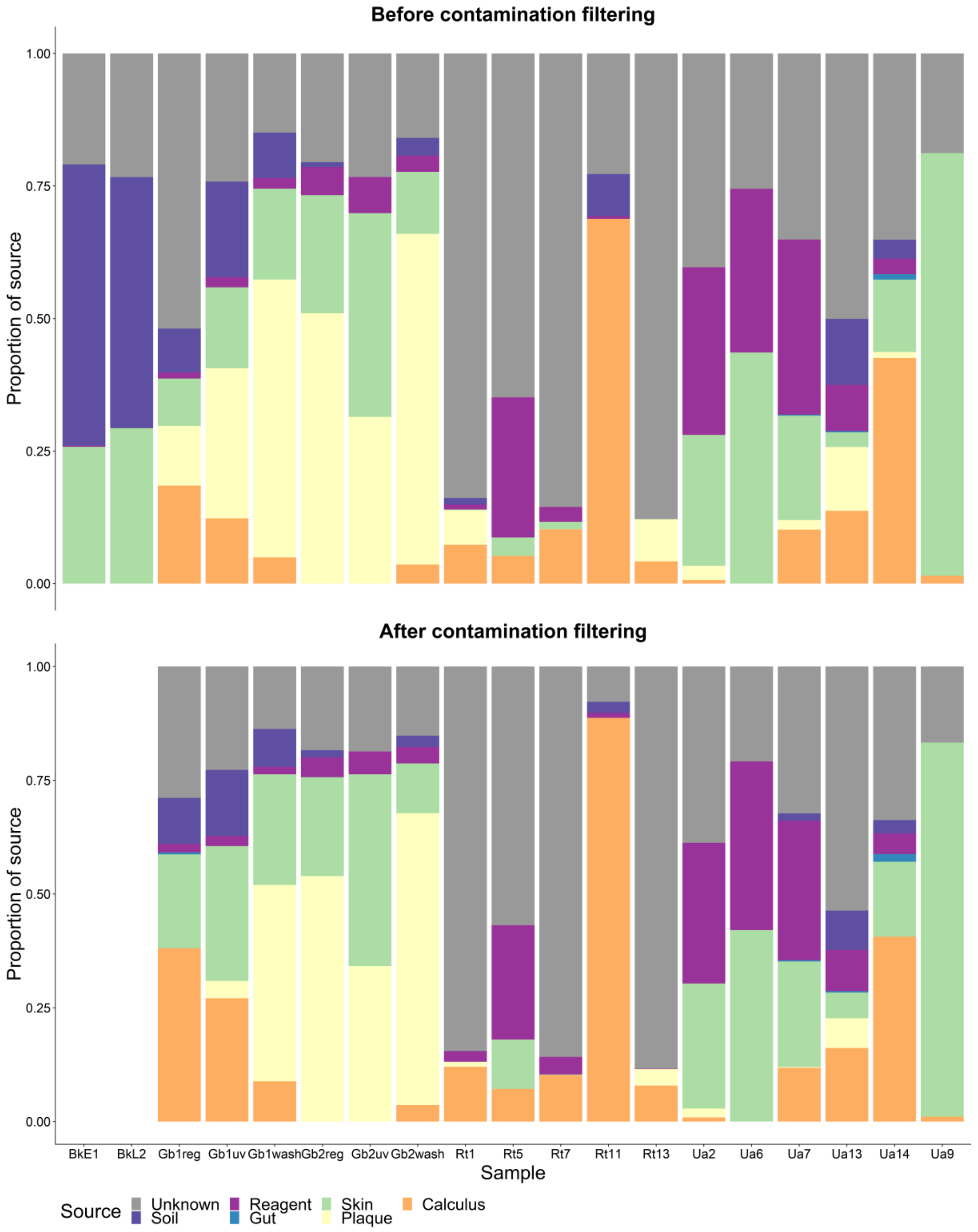


Supplementary Figure 1. Proportions of source contributions to the microbial communities (identified taxonomically at the species level) contained in the dental calculus samples estimated by SourceTracker, using modern human oral, human skin, human gut, laboratory reagent, and soil microbiome datasets as sources. Top: Before filtering out contaminant taxa. Bottom: After filtering of taxa based on the described filtering procedure (fragment length, presence in blanks and relative abundance). Note the proportional reduction of the "soil" source in the samples after filtering.

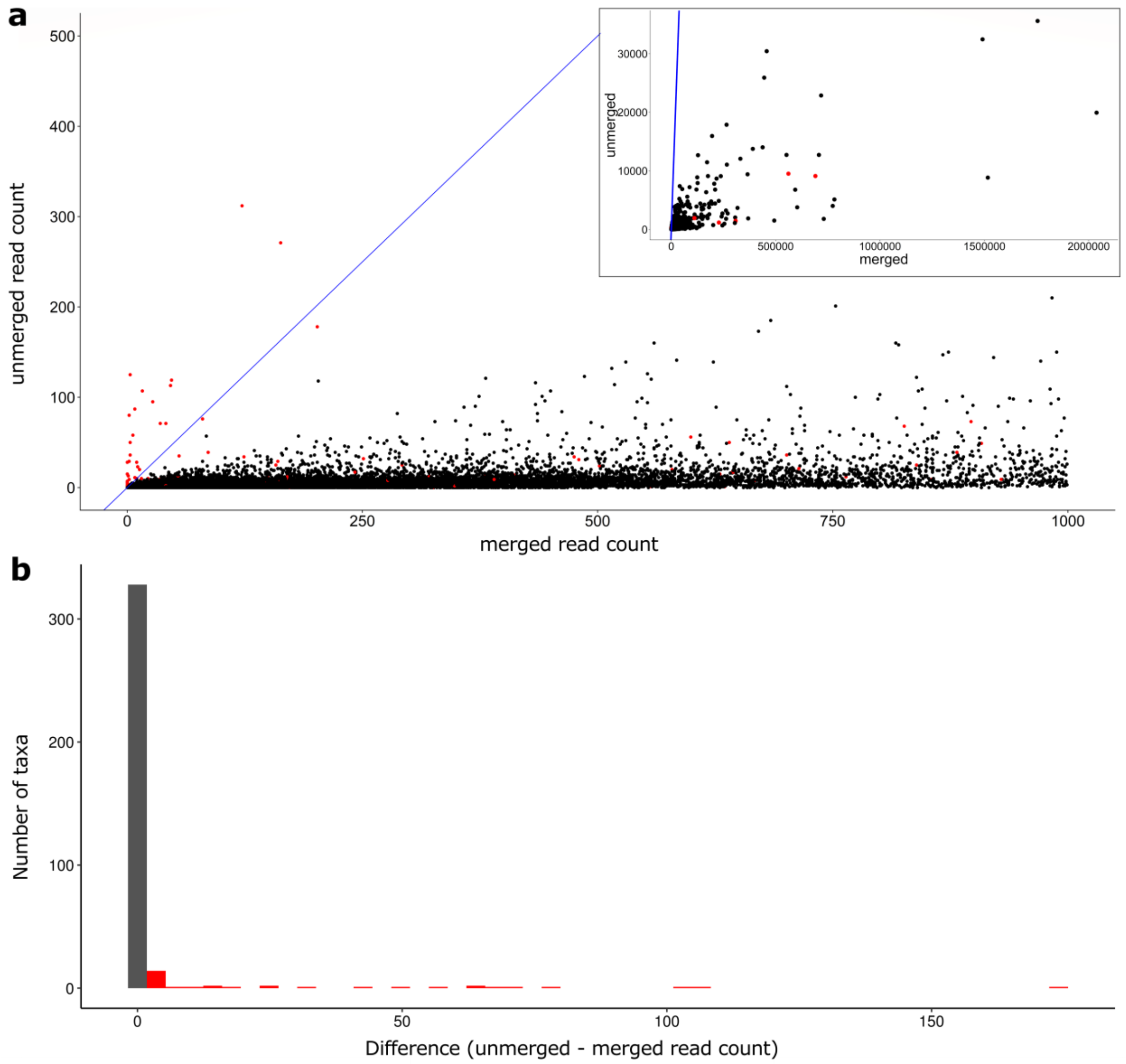


Supplementary Figure 2. Identifying potential contaminant taxa from merged and unmerged reads. **a)** Comparison of raw read counts between merged and unmerged reads, where each point represents a single taxon in a single sample. Taxa above the blue parity line (y=x) were more abundant in the unmerged reads in at least one sample, and were investigated further as possible modern contaminants (see Materials and Methods). These potential contaminants are highlighted in red for all samples, including those in which a possible contaminant fell below the parity line. Note that this is a zoomed-in plot to allow visualisation of the pattern, the full plot is shown in the inset, where it is clear that the vast majority of taxa within all samples are more abundant in the merged reads. **b)** Distribution of the differences in read number, calculated as unmerged minus merged reads, only shown for differences > 0 (i.e. those falling above the blue parity line in (a)). Taxa identified as outliers (defined as a difference > 1.5 interquartile ranges above the third), shown in red. These were filtered out as putative contaminants if observed in at least one sample.

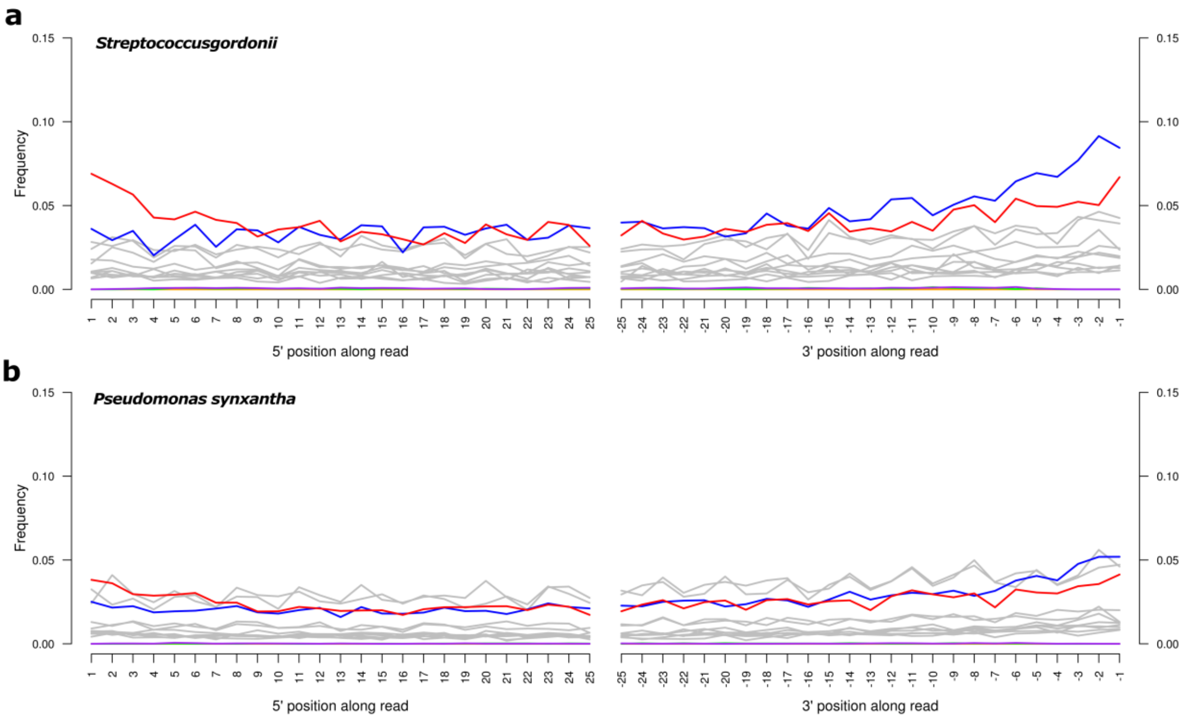


Supplementary Figure 3. Example patterns of post-mortem DNA damage for **(a)** *Streptococcus gordonii*, an authentic oral bacteria; and **(b)** *Pseudomonas synxantha*, a soil bacterium and likely modern contaminant that was removed from the dataset using the described filtering procedure. Deamination rates shown here are from sample Rt7. Red: C-to-T substitution, blue: G-to-A substitutions, grey: all other substitutions.

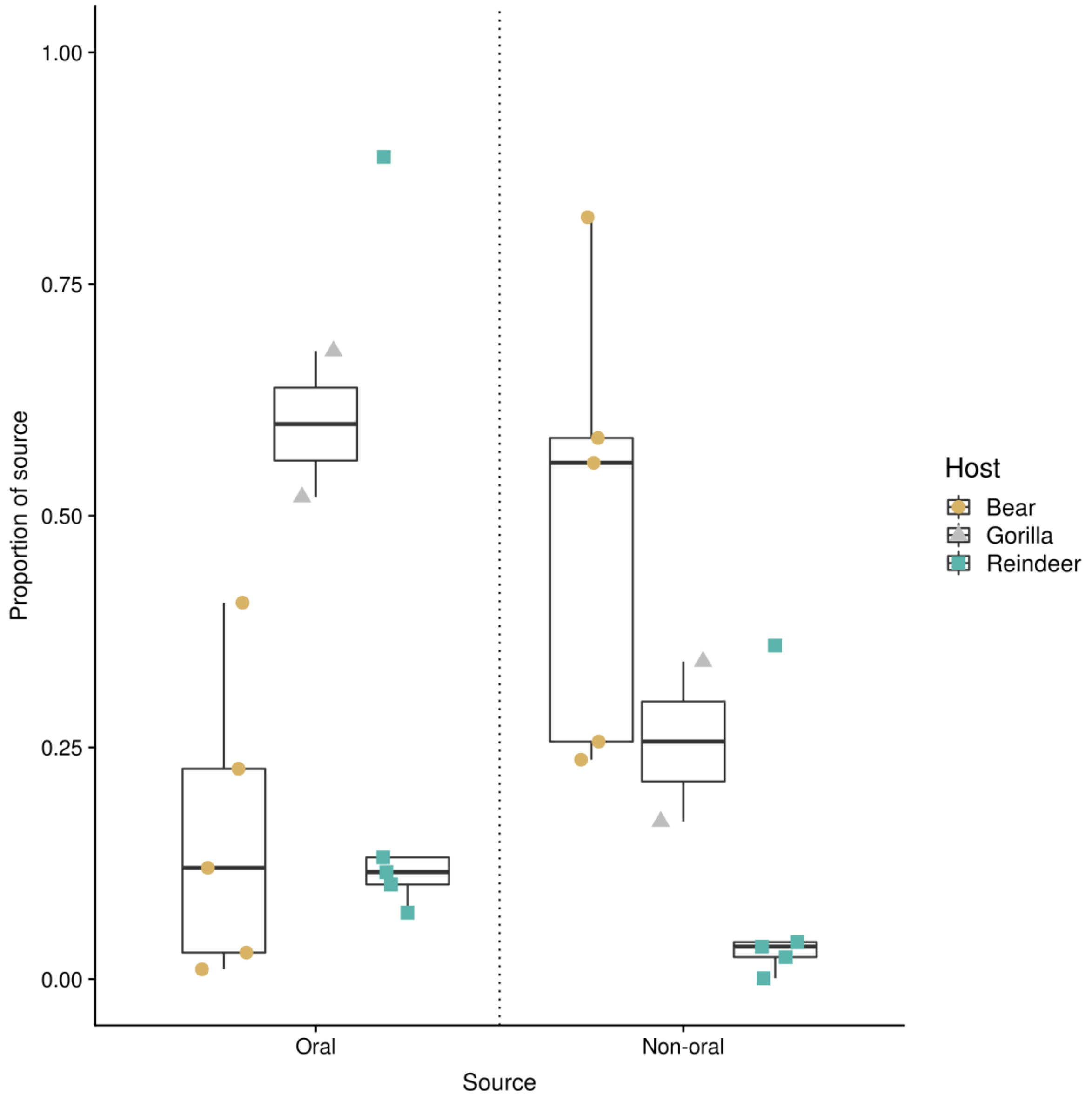


Supplementary Figure 4. Proportions (visualised as Tukey boxplots) of microbial taxa assigned by SourceTracker to oral and non-oral microbial taxa in bear (black), gorilla (dark grey) and reindeer (light grey) dental calculus samples. Oral taxa include those assigned to human historical dental calculus and modern dental plaque microbiomes; non-oral taxa include those assigned to human skin, human gut, soil and laboratory microbiome sources.

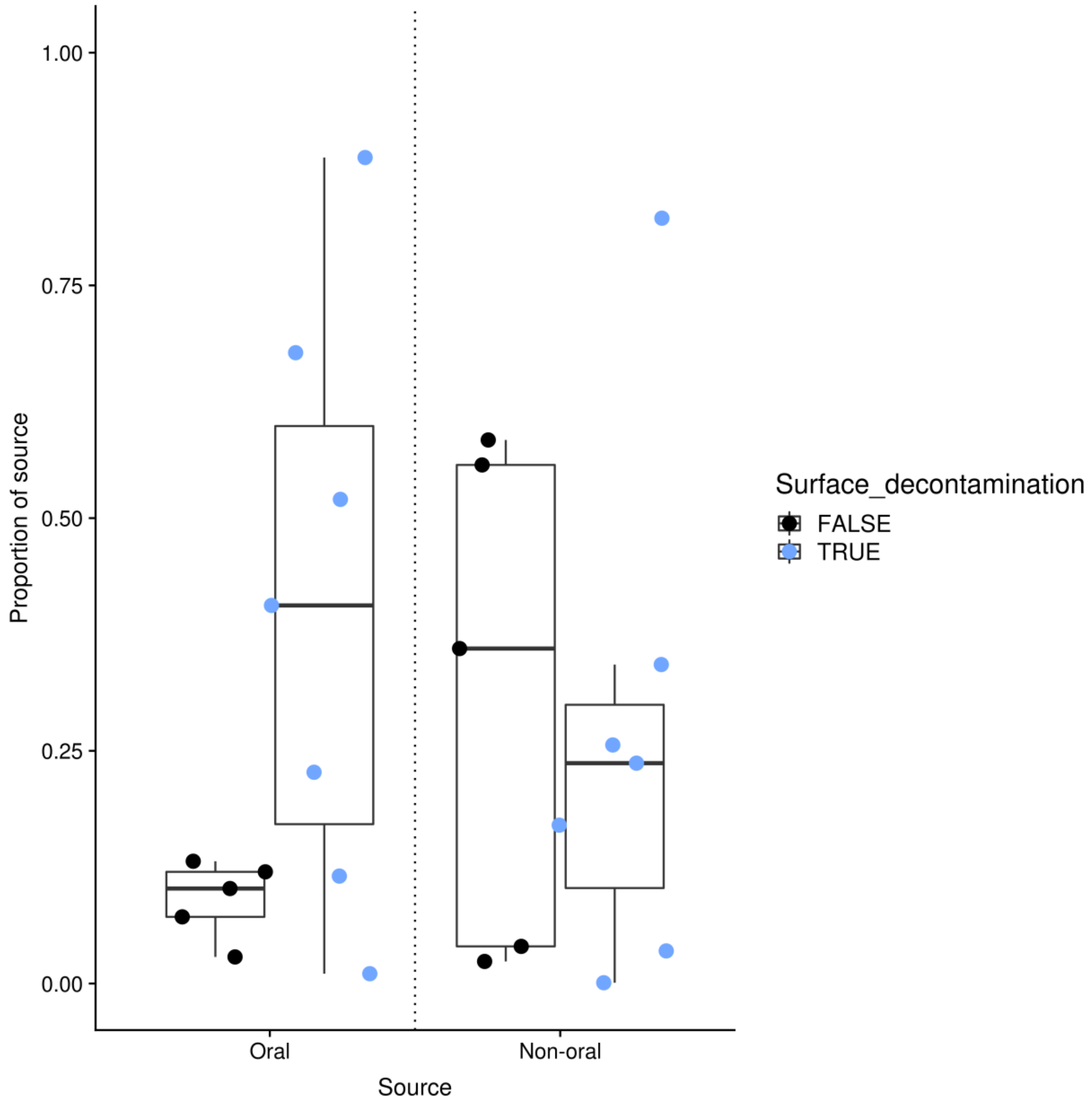


Supplementary Figure 5. Proportions (visualised as Tukey boxplots) of microbial taxa assigned by SourceTracker to oral and non-oral microbial taxa in samples that were surface decontaminated before DNA extraction (grey) and those that were extracted without surface decontamination (black). Surface decontamination consisted of a 10 min UV exposure and/or a 30 sec wash in 0.5M EDTA (Table S1). Oral taxa include those assigned to human historical dental calculus and modern dental plaque microbiomes; non-oral taxa include those assigned to human skin, human gut, soil and laboratory microbiome sources.

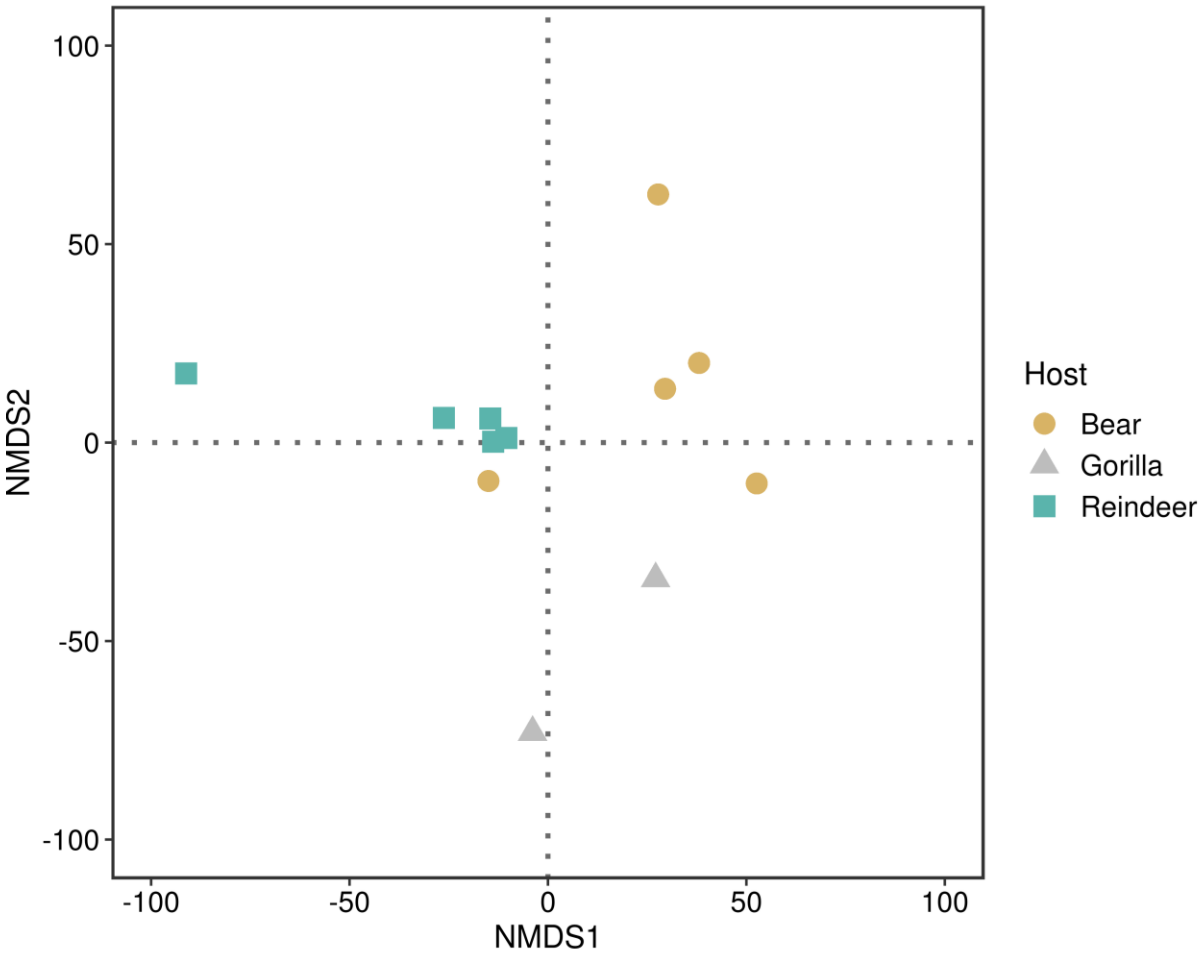


Supplementary Figure 6. Beta diversity based on the CLR normalised microbial abundance data. Non-metric multidimensional scaling (NMDS) using a distance matrix based on the Euclidean distance. Samples are coloured by host species.

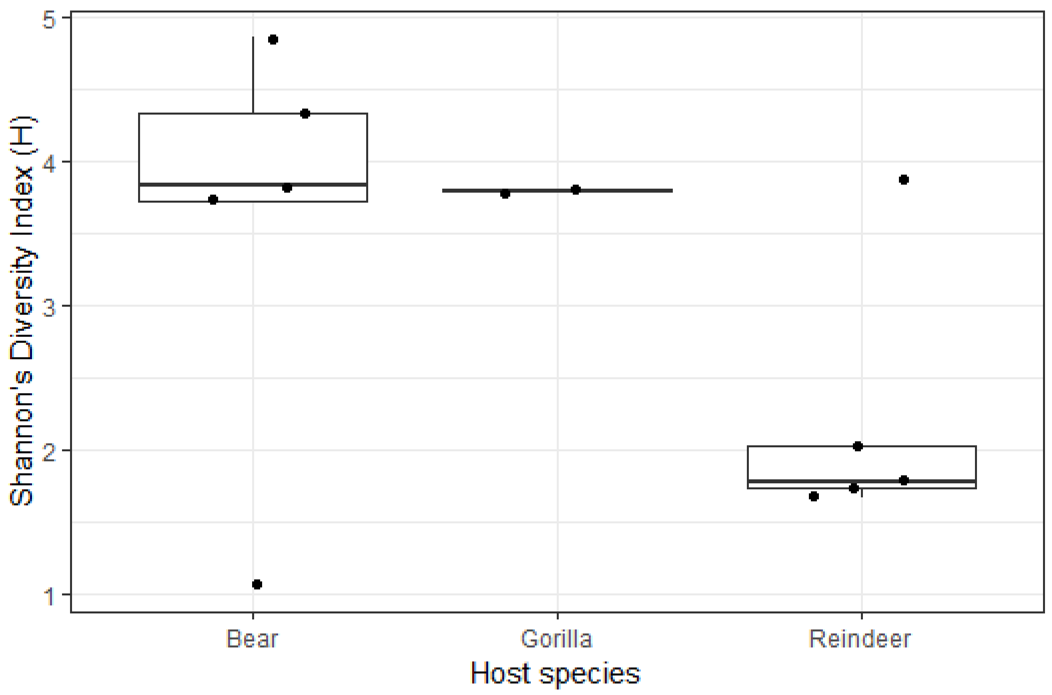


Supplementary Figure 7. Alpha diversity (visualised as Tukey boxplots) in samples from the three host species measured as the Shannon diversity index. Reindeer show overall lower diversity values.

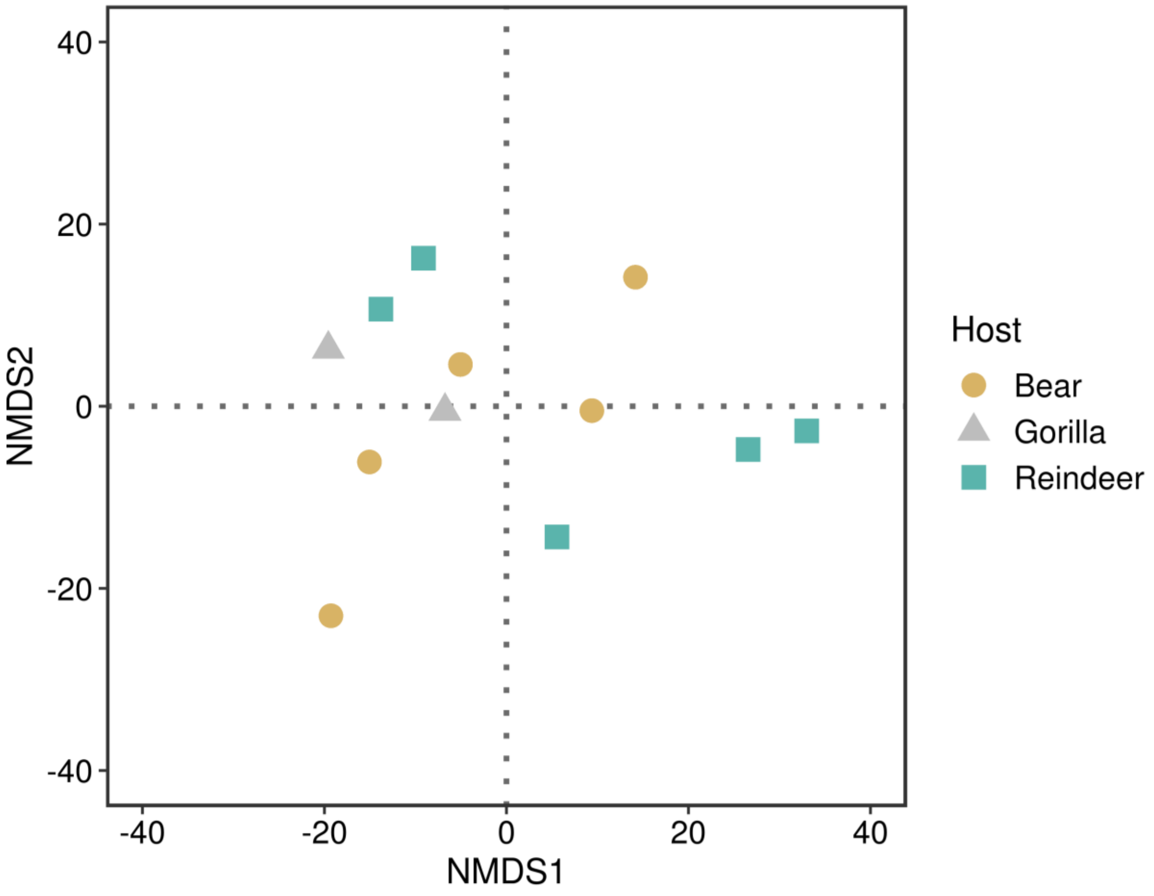


Supplementary Figure 8. Beta diversity based on the CLR normalised metabolic pathways abundance data from the HumanN2 functional analysis. Non-metric multidimensional scaling (NMDS) using a distance matrix based on the Euclidean distance. Samples are coloured by host species.

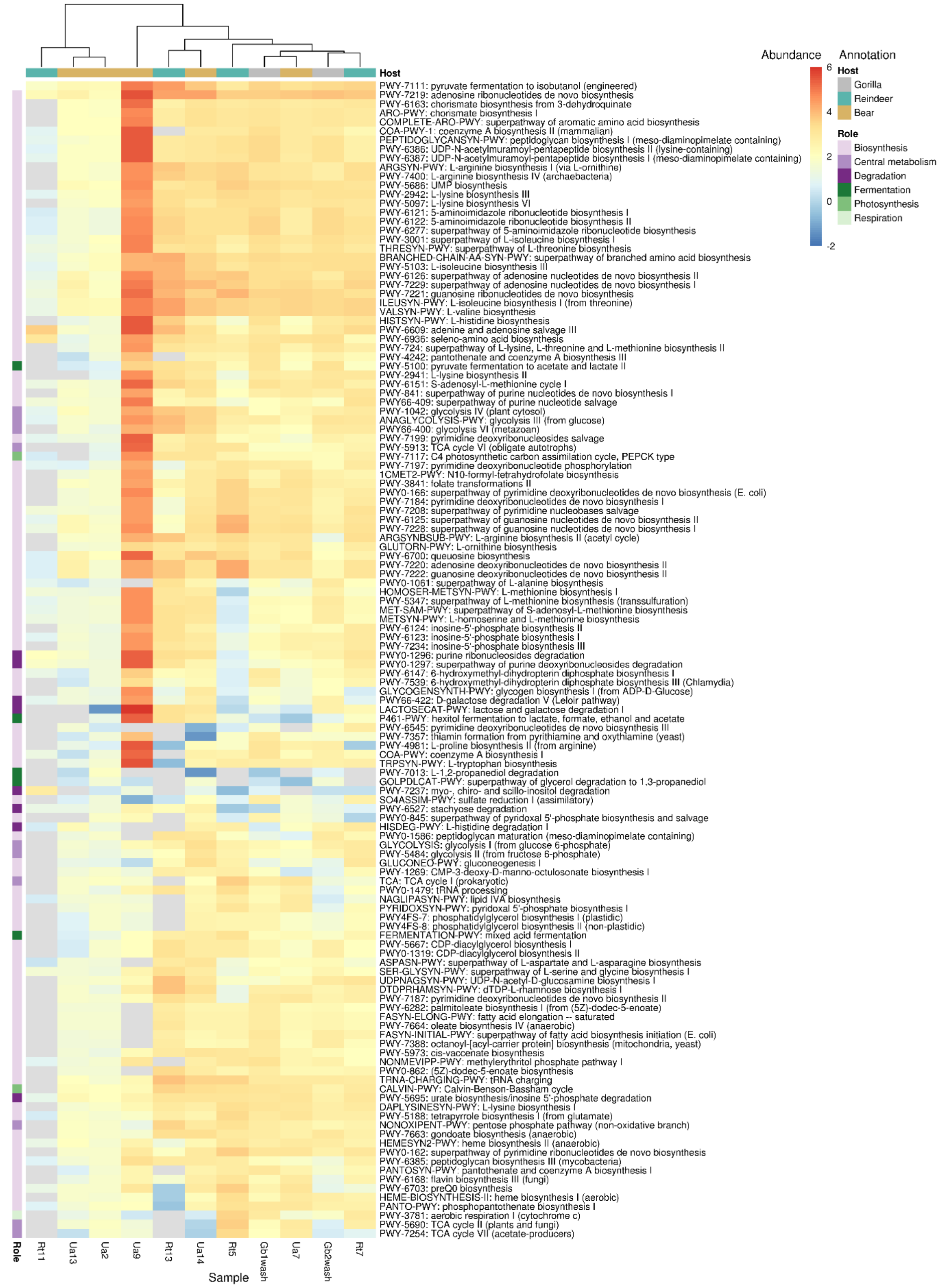


Supplementary Figure 9. Core functions are shared by the oral microbiome of gorillas, bears and reindeer. CLR normalised abundance of MetaCyc metabolic pathways containing >50% of required enzymes, as identified in by HumanN2 in each study sample. Taxa that were not detected in a sample are coloured grey. Metabolic pathways are annotated with their main role in metabolism. Samples and pathways are hierarchically clustered by dissimilarity implemented by pheatmap.

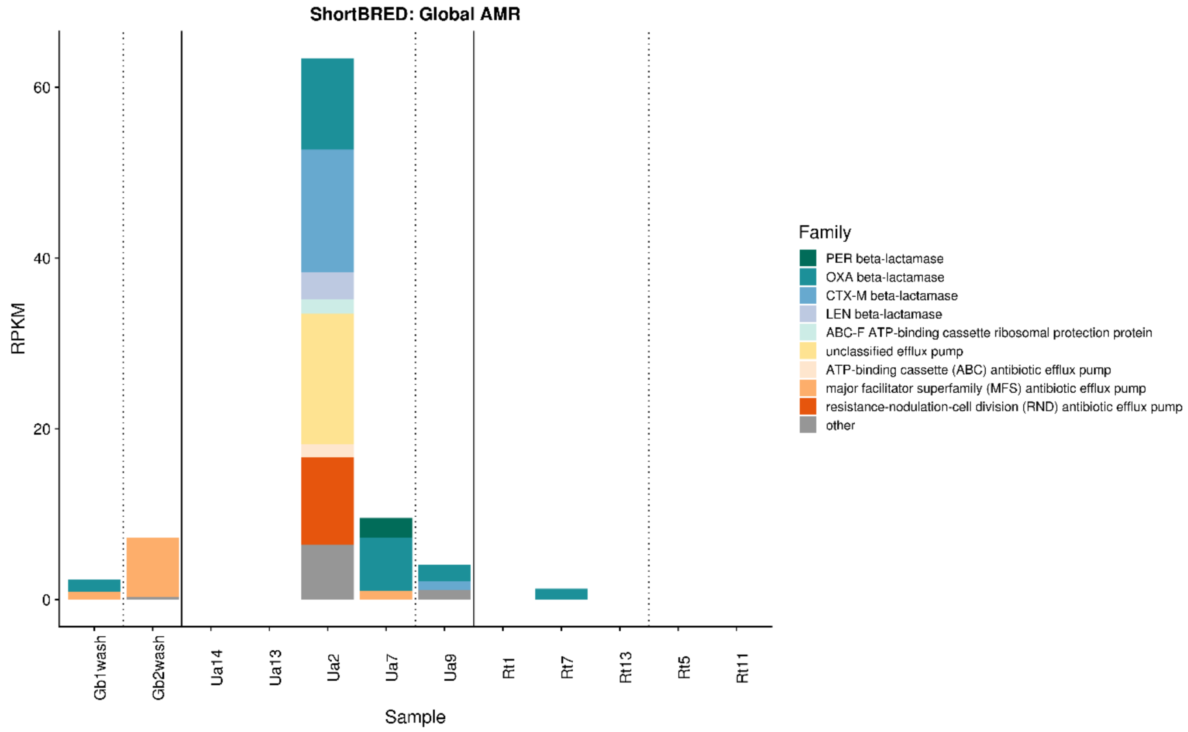


Supplementary Figure 10. AMR genes recovered in historical dental calculus specimens by the marker-based tool ShortBRED. Normalised counts (reads per kilobase of reference sequence per million sample reads [RPKM]) for each family in each sample are displayed. Samples are grouped by host species and ordered by year with pre-1940 samples separated from post-1940 samples by dashed vertical lines. The top nine most abundant AMR gene families are shown, with the remainder grouped into ‘other’.

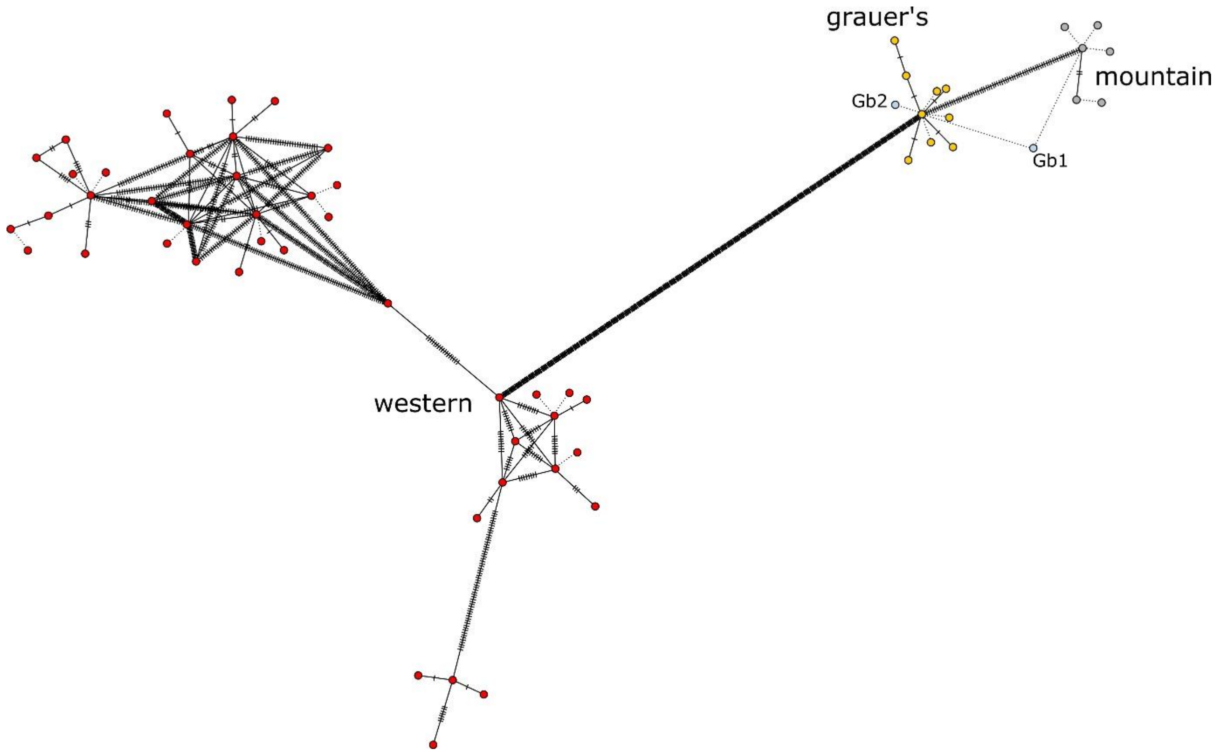


Supplementary Figure 11. mtDNA haplotype network for gorillas. Each dot represents a sample, with ticks on the connecting lines showing the number of base pair substitutions between the haplotypes. Dotted lines represent identical haplotypes or in the case of dental calculus samples (in light blue), the predicted most closely related haplotype. Gb2 clusters with the Grauer's gorillas, consistent with its museum-recorded subspecies designation. Gb1, a museum-designated mountain gorilla, groups with the Eastern gorillas but based on solely mtDNA cannot be identified at the subspecies level. However, using whole genome data, Gb1 clusters more closely to mountain gorillas (Figure 5B).

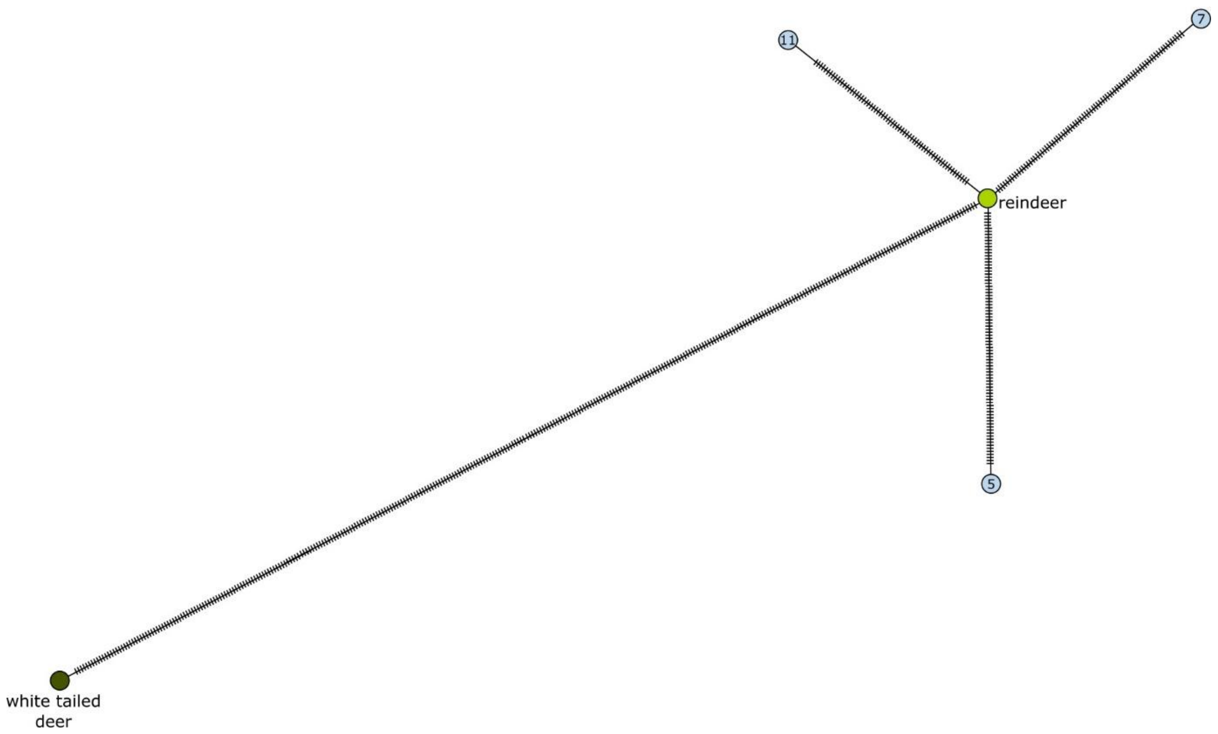


Supplementary Figure 12. mtDNA haplotype network for reindeer. Each dot represents a sample, with ticks on the connecting lines showing the number of base pair substitutions between the haplotypes. Genomic profiles recovered from dental calculus samples are shown in light blue. Only three dental calculus samples contained enough host mtDNA to allow haplotype prediction. As only one reindeer reference genome is currently available, no sub-species or population-level inferences could made.

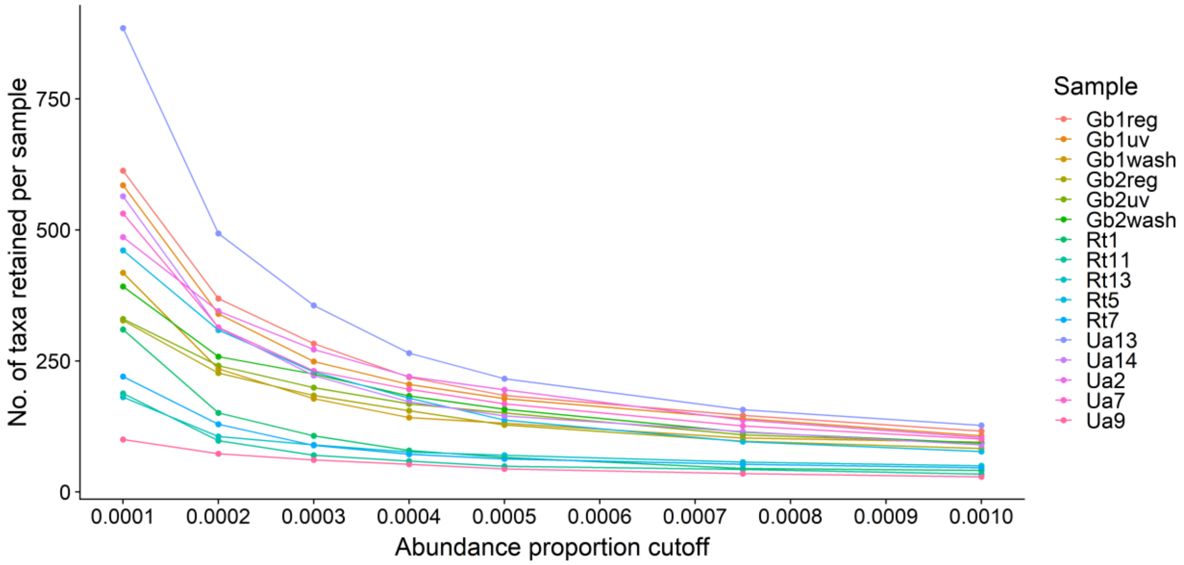


Supplementary Figure 13. Investigation of abundance filtering thresholds on community complexity, based on microbial relative abundance. The number of taxa retained in each sample (coloured lines) at each threshold is shown.

**Supplementary Tables**

Supplementary Table 1. Sample metadata, see Supplementary Data.

Supplementary Table 2. PERMANOVA results from microbial analyses.

| Dataset | Distance matrix | Variable | Degrees of freedom | Sequential sum of squares | Mean squares | F statistic | Partial R-squared value | *p* value |
|---|---|---|---|---|---|---|---|---|
| Taxonomic presence/ absence | Jaccard | Host species[1] | 2 | 1.0443 | 0.5222 | 1.8778 | 0.2967 | **0.0060** |
| | | Decontamination | 1 | 0.3356 | 0.3356 | 1.2067 | 0.0953 | 0.1902 |
| | | No. of reads | 1 | 0.2210 | 0.2210 | 0.7946 | 0.0628 | 0.7696 |
| | | Proportion of human DNA | 1 | 0.2502 | 0.2503 | 0.8999 | 0.0711 | 0.5882 |
| | | Residuals | 6 | 1.6685 | 0.2781 | | 0.4741 | |
| Taxonomic abundance, CLR normalised | Euclidean | Host species | 2 | 20077 | 10038 | 2.2693 | 0.3163 | **< 0.001** |
| | | Decontamination | 1 | 5998 | 5998 | 1.3559 | 0.0945 | 0.0997 |
| | | No. of reads | 1 | 5822 | 5822 | 1.3162 | 0.0917 | 0.1611 |
| | | Proportion of human DNA | 1 | 5039 | 5039 | 1.1392 | 0.0794 | 0.3034 |
| | | Residuals | 6 | 26541 | 4424 | | 0.4181 | |
| Metabolic pathway abundance, CLR normalised | Euclidean | Host species | 2 | 1852 | 926 | 1.3519 | 0.2177 | 0.1550 |
| | | Decontamination | 1 | 718 | 718 | 1.0481 | 0.0844 | 0.3412 |
| | | No. of reads | 1 | 1141 | 1141 | 1.6659 | 0.1342 | 0.1000 |
| | | Residuals | 7 | 4795 | 685 | | 0.5637 | |

[1] A distance-based test for homogeneity of multivariate dispersions did not detect differences in group homogeneity between the study host species ($F_{2,9} = 0.9447$, $p = 0.4865$). Average distances from median for bear, gorilla, and reindeer samples were, respectively, 0.4426, 0.3372, 0.4824.

Supplementary Table 3. Oral taxa unique to host species.

| Host species | No. of taxa | Unique taxa | Unique taxa shared by >50% of samples | Likely origin[1] of microbial taxa | | |
|---|---|---|---|---|---|---|
| | | | | Oral | Host-associated | Other |
| Gorilla[2] | 304 | 74 | 0 | 28.40% | 18.90% | 52.70% |
| Bear | 547 | 257 | 29 | 37.90% | 51.70% | 10.40% |
| Reindeer | 372 | 175 | 5 | 0.00% | 60.00% | 40.00% |

[1] Microbial taxa were classified as oral, non-oral but host-associated in mammals, and not associated with mammals, based on literature searches.

[2] Percentages for gorillas were calculated based on the number of unique taxa, whereas for reindeer and bear percentages were calculated based on the number of unique taxa shared by >50% of samples within a host.

Supplementary Table 4. Metagenome assembled genome (MAG) taxonomy and quality estimates.

| Sample | Taxonomy | Completeness (%) | Contamination (%) | Mean coverage per MAG site | Closest reference genome[1] | Damage (C->T %) at 5' end (terminal, penultimate bases)[2] | Proportion of reads identified by Kraken[3] |
|---|---|---|---|---|---|---|---|
| Ua9 | *Lactobacillus C paracasei* | 96.13 | 0.03 | 34.3 | GCF_000829035.1 | 2.5, 4.1 | 0.057 |
| Ua9 | *Streptococcus ratti* | 96.59 | 0.59 | 116.8 | GCF_000286075.1 | 2.7, 4.3 | 0.872 |
| Ua14 | *Petrimonas sp.* | 94.2 | 1.64 | 23.1 | NA | NA | 0.00376 |
| Ua14 | *Streptococcus orisasini* | 92.46 | 2.75 | 14.5 | GCF_001431045.1 | 4.3, 4.0 | 0.301 |
| Ua14 | *'UBA11524' sp. (Christensenellales)* | 91.76 | 2.42 | 15.8 | NA | NA | NA |
| Rt7 | *Streptococcus sp.* | 97.73 | 0.37 | 69.9 | GCF_001642085.1 (*Streptococcus pantholopis*) | 6.5, 6.7 | 0.0813 |
| Rt13 | *Streptococcus sp.* | 91.48 | 0.59 | 39.3 | GCF_001642085.1 (*Streptococcus pantholopis*) | 8.4, 8.7 | 0.0658 |
| Rt13 | *Haemophilus B sp.* | 91.39 | 1.59 | 28.8 | GCF_001647695.1 (*Haemophilus ducreyi*) | 6.5, 6.2 | 0.00599 |

[1] As identified by fastANI during GTDB-TK taxonomy analysis; fastANI failed for two MAGs and novel taxonomy was determined using relative evolutionary divergence of the MAG within the reference tree.

[2] Damage data based on mapDamage comparison of reads mapped to closest genome reference.

[3] Proportion of reads identified at genus-level (or species-level in the case of *L. paracasei*) out of total number of reads classified by Kraken at the genus and species levels.

Supplementary Table 5. Host DNA recovery statistics.

| Sample ID | Host | No. of host reads | Nuclear DNA | | Mitochondrial DNA | | Mitochondrial genome reconstruction | | | | | |
|---|---|---|---|---|---|---|---|---|---|---|---|---|
| | | | Mean coverage | % covered at ≥1x | Mean coverage | % covered at ≥1x | Coding length[1] (nt) | No. of reads used for assembly | Maximum coverage | Average total coverage | Average consensus quality | GC% |
| Gb1wash | Gorilla | 24931 | 0.001 | 0.056 | 0.511 | 39.08 | | | | | | |
| Gb2wash | Gorilla | 1227196 | 0.009 | 0.789 | 13.035 | 99.46 | 15449 | 2700 | 29 | 13.65 | 87 | 43.97 |
| Ua2 | Brown bear | 218318 | 0.004 | 0.346 | 3.354 | 92.22 | | | | | | |
| Ua6 | Brown bear | 3941520 | 0.077 | 6.952 | 16.527 | 96.40 | 15443 | 4174 | 40 | 19.39 | 89 | 41.17 |
| Ua7 | Brown bear | 19104 | 0.001 | 0.040 | 0.227 | 15.44 | | | | | | |
| Ua13 | Brown bear | 41004 | 0.001 | 0.076 | 0.157 | 14.96 | | | | | | |
| Ua14 | Brown bear | 6428823 | 0.191 | 16.503 | 92.563 | 97.96 | 15443 | 19196 | 150 | 95.51 | 90 | 41.11 |
| Ua9 | Brown bear | 4690 | 0.000 | 0.007 | 0.131 | 13.27 | | | | | | |
| Rt1 | Reindeer | 3617 | 0.000 | 0.004 | 0.026 | 2.58 | | | | | | |
| Rt5 | Reindeer | 1653467 | 0.025 | 2.446 | 8.267 | 97.65 | 15482 | 2045 | 24 | 10.93 | 88 | 36.26 |
| Rt7 | Reindeer | 730021 | 0.011 | 1.071 | 2.352 | 91.21 | | | | | | |
| Rt11 | Reindeer | 24838941 | 0.308 | 21.261 | 173.890 | 99.96 | 15482 | 31958 | 240 | 155.41 | 90 | 36.29 |
| Rt13 | Reindeer | 27916 | 0.000 | 0.020 | 0.066 | 5.62 | | | | | | |

[1] Length of covered coding region, excluding the hypervariable control region (as it could not be reconstructed in most samples).

Supplementary Table 6. Sequencing adapter and real-time PCR assay primer sequences.

| Type | Sequence (5'-3')[1] |
|---|---|
| Barcoded P5 forward adapter | CTTTCCCTACACGACGCTCTTCCGATCTxxxxxxx |
| Barcoded P7 forward adapter | GTGACTGGAGTTCAGACGTGTGCTCTTCCGATCTxxxxxxx |
| Barcoded P5/P7 reverse adapter | xxxxxxxAGATCG |
| Indexed P5 adapter primer | AATGATACGGCGACCACCGAGATCTACACxxxxxxxACACTCTTTCCCTACACGACGCTCTT |
| Indexed P7 adapter primer | CAAGCAGAAGACGGCATACGAGATxxxxxxxGTGACTGGAGTTCAGACGTGT |
| preHyb forward primer | CTTTCCCTACACGACGCTCTTC |
| preHyb reverse primer | GTGACTGGAGTTCAGACGTGTGCT |
| IS5 P5 primer | AATGATACGGCGACCACCGA |
| IS6 P7 primer | CAAGCAGAAGACGGCATACGA |

[1] For sample-specific barcode and adapter sequences (indicated by ‘x’ strings), see Supplementary Table 1 (Supplementary Data).

Supplementary Table 7. Samples used as sources for SourceTracker analysis.

| **Source environment** | **Citation** | **Accession numbers** |
|---|---|---|
| Human oral (modern supragingival plaque) | [1,2] | SRS015650<br>SRS018665<br>SRS018975<br>SRS019387<br>SRS023538 |
| Human oral (medieval dental calculus) | [3] | SRR6877300<br>SRR6877340<br>SRR6877350<br>SRR6877370<br>SRR6877399 |
| Human skin (palm) | [4] | SRS727524<br>SRS728264<br>SRS728285<br>SRS728312<br>SRS728355 |
| Human gut (stool) | [1,2] | SRS011271<br>SRS018656<br>SRS019397<br>SRS023526<br>SRS051031 |
| Soil (Alaskan topsoil permafrost) | [5] | ERR1017187<br>ERR1019366<br>ERR1022687<br>ERR1034454<br>ERR1035437 |
| Laboratory reagents (blank controls) | [6] | ERR584320<br>ERR584333<br>ERR584341<br>ERR584348 |

Supplementary Table 8. GenBank accessions of taxa identified and removed as contaminants, see Supplementary Data.

Supplementary Table 9. Investigation into post-mortem deamination and barcoded adapter bias, see Supplementary Data.

Supplementary Table 10. Summary of investigation into post-mortem deamination and barcoded adapter bias.

| | **Deamination drop at terminal base** | |
|---|---|---|
| **Barcode terminal base** | No | Yes |
| A/T at end | 32 | 6 |
| C/G at end | 6 | 30 |
